# Largest genome-wide association study for PTSD identifies genetic risk loci in European and African ancestries and implicates novel biological pathways

**DOI:** 10.1101/458562

**Authors:** Caroline M. Nievergelt, Adam X. Maihofer, Torsten Klengel, Elizabeth G. Atkinson, Chia-Yen Chen, Karmel W. Choi, Jonathan R.I. Coleman, Shareefa Dalvie, Laramie E. Duncan, Mark W. Logue, Allison C. Provost, Andrew Ratanatharathorn, Murray B. Stein, Katy Torres, Allison E. Aiello, Lynn M. Almli, Ananda B. Amstadter, Søren B Andersen, Ole A. Andreassen, Paul A. Arbisi, Allison E. Ashley-Koch, S. Bryn Austin, Esmina Avdibegovic, Dragan Babić, Marie Bækvad-Hansen, Dewleen G. Baker, Jean C. Beckham, Laura J. Bierut, Jonathan I. Bisson, Marco P. Boks, Elizabeth A. Bolger, Anders D. Børglum, Bekh Bradley, Megan Brashear, Gerome Breen, Richard A. Bryant, Angela C. Bustamante, Jonas Bybjerg-Grauholm, Joseph R. Calabrese, José M. Caldas-de-Almeida, Anders M. Dale, Mark J. Daly, Nikolaos P. Daskalakis, Jürgen Deckert, Douglas L. Delahanty, Michelle F. Dennis, Seth G. Disner, Katharina Domschke, Alma Dzubur-Kulenovic, Christopher R. Erbes, Alexandra Evans, Lindsay A. Farrer, Norah C. Feeny, Janine D. Flory, David Forbes, Carol E. Franz, Sandro Galea, Melanie E. Garrett, Bizu Gelaye, Joel Gelernter, Elbert Geuze, Charles Gillespie, Aferdita Goci Uka, Scott D. Gordon, Guia Guffanti, Rasha Hammamieh, Supriya Harnal, Michael A. Hauser, Andrew C. Heath, Sian M.J. Hemmings, David Michael Hougaard, Miro Jakovljevic, Marti Jett, Eric Otto Johnson, Ian Jones, Tanja Jovanovic, Xue-Jun Qin, Angela G. Junglen, Karen-Inge Karstoft, Milissa L. Kaufman, Ronald C. Kessler, Alaptagin Khan, Nathan A. Kimbrel, Anthony P. King, Nastassja Koen, Henry R. Kranzler, William S. Kremen, Bruce R. Lawford, Lauren A.M. Lebois, Catrin E. Lewis, Sarah D. Linnstaedt, Adriana Lori, Bozo Lugonja, Jurjen J. Luykx, Michael J. Lyons, Jessica Maples-Keller, Charles Marmar, Alicia R. Martin, Nicholas G. Martin, Douglas Maurer, Matig R. Mavissakalian, Alexander McFarlane, Regina E. McGlinchey, Katie A. McLaughlin, Samuel A. McLean, Sarah McLeay, Divya Mehta, William P. Milberg, Mark W. Miller, Rajendra A. Morey, Charles Phillip Morris, Ole Mors, Preben B. Mortensen, Benjamin M. Neale, Elliot C. Nelson, Merete Nordentoft, Sonya B. Norman, Meaghan O’Donnell, Holly K. Orcutt, Matthew S. Panizzon, Edward S. Peters, Alan L. Peterson, Matthew Peverill, Robert H. Pietrzak, Melissa A. Polusny, John P. Rice, Stephan Ripke, Victoria B. Risbrough, Andrea L. Roberts, Alex O. Rothbaum, Barbara O. Rothbaum, Peter Roy-Byrne, Ken Ruggiero, Ariane Rung, Bart P. F. Rutten, Nancy L. Saccone, Sixto E. Sanchez, Dick Schijven, Soraya Seedat, Antonia V. Seligowski, Julia S. Seng, Christina M. Sheerin, Derrick Silove, Alicia K. Smith, Jordan W. Smoller, Nadia Solovieff, Scott R. Sponheim, Dan J. Stein, Jennifer A. Sumner, Martin H. Teicher, Wesley K. Thompson, Edward Trapido, Monica Uddin, Robert J. Ursano, Leigh Luella van den Heuvel, Miranda van Hooff, Eric Vermetten, Christiaan H. Vinkers, Joanne Voisey, Yunpeng Wang, Zhewu Wang, Thomas Werge, Michelle A. Williams, Douglas E. Williamson, Sherry Winternitz, Christiane Wolf, Erika J. Wolf, Jonathan D. Wolff, Rachel Yehuda, Keith A. Young, Ross McD. Young, Hongyu Zhao, Lori A. Zoellner, Israel Liberzon, Kerry J. Ressler, Magali Haas, Karestan C. Koenen

**Affiliations:** University of California San Diego, Department of Psychiatry, La Jolla, CA, US.; Veterans Affairs San Diego Healthcare System, Center of Excellence for Stress and Mental Health, San Diego, CA, US.; Veterans Affairs San Diego Healthcare System, Research Service, San Diego, CA, US; Harvard Medical School, Department of Psychiatry, Boston, MA, US; McLean Hospital, Belmont, MA, US; Broad Institute, Stanley Center for Psychiatric Research, Cambridge, MA, US; Massachusetts General Hospital, Analytic and Translational Genetics Unit, Boston, MA, US; Massachusetts General Hospital, Psychiatric and Neurodevelopmental Genetics Unit (PNGU), Boston, MA, US; Harvard T.H, Chan School of Public Health, Department of Epidemiology, Boston, MA, US; Massachusetts General Hospital, Department of Psychiatry, Boston, MA, US; King’s College London, Social, Genetic and Developmental Psychiatry Centre, Institute of Psychiatry, Psychology and Neuroscience, London, GB; King’s College London, NIHR BRC at the Maudsley, London, GB; University of Cape Town, SA MRC Unit on Risk & Resilience in Mental Disorders, Department of Psychiatry, Cape Town, Western Cape, ZA.; Stanford University, Department of Psychiatry and Behavioral Sciences, Stanford, CA, US; VA Boston Healthcare System, National Center for PTSD, Boston, MA, US; TH Chan School of Public Health, Department of Epidemiology, Boston, MA, US; Veterans Affairs San Diego Healthcare System, Psychiatry Service, San Diego, CA, US; University of North Carolina at Chapel Hill, Department of Epidemiology, Chapel Hill, NC, US; Carter Consulting, Atlanta, GA, US; Virginia Institute for Psychiatric and Behavioral Genetics, Department of Psychiatry, Richmond, VA, US; The Danish Veteran Centre, Research and Knowledge Centre, Ringsted, Sjaelland, DK; University of Oslo, Institute of Clinical Medicine, Oslo, NO.; Minneapolis VA Health Care System, Mental Health Service Line, Minneapolis, MN, US.; Duke University, Department of Psychiatry, Durham, NC, US.; Boston Children’s Hospital, Division of Adolescent and Young Adult Medicine, Boston, MA, US.; Brigham and Women’s Hospital, Channing Division of Network Medicine, Boston, MA, US.; Harvard School of Public Health, Department of Social and Behavioral Sciences, Boston, MA, US.; University Clinical Center of Tuzla, Department of Psychiatry, Tuzla, BA.; University Clinical Center of Mostar, Department of Psychiatry, Mostar, BA.; Statens Serum Institut, Department for Congenital Disorders, Copenhagen, DK.; The Lundbeck Foundation Initiative for Integrative Psychiatric Research, iPSYCH, DK.; Duke University, Department of Psychiatry and Behavioral Sciences, Durham, NC, US.; Durham VA Medical Center, Research, Durham, NC, US.; VA Mid-Atlantic Mental Illness Research, Education, and Clinical Center (MIRECC), Genetics Research Laboratory, Durham, NC, US.; Washington University in Saint Louis School of Medicine, Department of Psychiatry, Saint Louis, MO, US.; Cardiff University, National Centre for Mental Health, MRC Centre for Psychiatric Genetics and Genomics, Cardiff, South Glamorgan, GB.; UMC Utrecht Brain Center Rudolf Magnus, Department of Translational Neuroscience, Utrecht, Utrecht, NL.; Aarhus University, Centre for Integrative Sequencing, iSEQ, Aarhus, DK.; Aarhus University, Department of Biomedicine - Human Genetics, Aarhus, DK; Atlanta VA Health Care System, Mental Health Service Line, Decatur, GA, US.; Emory University, Department of Psychiatry and Behavioral Sciences, Atlanta, GA, US.; Louisiana State University Health Sciences Center, School of Public Health and Department of Epidemiology, New Orleans, LA, US.; University of New South Wales, Department of Psychology, Sydney, NSW, AU.; University of Michigan Medical School, Division of Pulmonary and Critical Care Medicine, Department of Internal Medicine, Ann Arbor, Mi, US.; University Hospitals, Department of Psychiatry, Cleveland, OH, US.; CEDOC -Chronic Diseases Research Centre, Lisbon Institute of Global Mental Health, Lisbon, PT.; Unversity of California, San Diego, Department of Radiology, Department of Neurosciences, La Jolla, CA, US.; Cohen Veterans Bioscience, Cambridge, MA, US.; Icahn School of Medicine at Mount Sinai, Department of Psychiatry, New York, NY, US.; University Hospital of Würzburg, Center of Mental Health, Psychiatry, Psychosomatics and Psychotherapy, Würzburg, DE.; Kent State University, Department of Psychological Sciences, Kent, OH, US.; Kent State University, Research and Sponsored Programs, Kent, OH, US.; Duke University, Duke Molecular Physiology Institute, Durham, NC, US.; Minneapolis VA Health Care System, Research Service Line, Minneapolis, MN, US.; Medical Center-University of Freiburg, Faculty of Medicine, Department of Psychiatry and Psychotherapy, Freiburg, DE.; University Clinical Center of Sarajevo, Department of Psychiatry, Sarajevo, BA.; Minneapolis VA Health Care System, Center for Care Delivery and Outcomes Research (CCDOR), Minneapolis, MN, US.; University of Minnesota, Department of Psychiatry, Minneapolis, MN, US.; Boston University School of Medicine, Department of Medicine, Boston, MA, US.; Case Western Reserve University, Department of Psychological Sciences, Cleveland, OH, US.; University of Melbourne, Department of Psychiatry, Melbourne, VIC, AU.; Boston University, Department of Psychological and Brain Sciences, Boston, MA, US.; US Department of Veterans Affairs, Department of Psychiatry, West Haven, CT, US.; Yale University School of Medicine, Department of Genetics and Neuroscience, New Haven, CT, US.; Netherlands Ministry of Defence, Research Center Military Mental Healthcare, Utrecht, Utrecht, NL.; UMC Utrecht Brain Center Rudolf Magnus, Department of Psychiatry, Utrecht, Utrecht, NL.; University Clinical Centre of Kosovo, Department of Psychiatry, Prishtina, Kosovo, XK.; QIMR Berghofer Medical Research Institute, Department of Genetics and Computational Biology, Brisbane, Queensland, AU.; US Army Medical Research and Materiel Command, USACEHR, Fort Detrick, MD, US.; Broad Institute, Stanley Center for Psychiatric Research, Cambridge, MA, US.; Washington University in Saint Louis School of Medicine, Department of Genetics, Saint Louis, MO, US.; Stellenbosch University Faculty of Medicine and Health Sciences, Department of Psychiatry, Cape Town, Western Cape, ZA.; University Hospital Center of Zagreb, Department of Psychiatry, Zagreb, HR.; RTI International, Behavioral Health and Criminal Justice Division, Research Triangle Park, NC, US.; Kent State University, Department of Pschological Sciences, Kent, OH, US.; The Danish Veteran Center, Research and Knowledge Centre, Ringsted, Zealand, DK.; University of Copenhagen, Department of Psychology, Copenhagen, DK.; Harvard Medical School, Department of Health Care Policy, Boston, MA, US.; VA Mid-Atlantic Mental Illness Research, Education, and Clinical Center (MIRECC), Genetics Research Laboratory, Durham, North Carolina, US.; University of Michigan Medical School, Department of Psychiatry, Ann Arbor, MI, US.; University of Cape Town, SA MRC Unit on Risk & Resilience in Mental Disorders, Department of Psychiatry, Cape Town, Observatory, Western Cape, ZA.; University of Pennsylvania Perelman School of Medicine, Department of Psychiatry, Philadelphia, PA, US.; University of California San Diego, Department of Psychiatry and Department of Family Medicine and Public Health, La Jolla, CA, US.; Queensland University of Technology, IHBI, Brisbane, QLD, AU.; Queensland University of Technology, School of Biomedical Sciences, Brisbane, QLD, AU.; UNC Institute for Trauma Recovery, Department of Anesthesiology, Chapel Hill, NC, US.; Emory University, Department of Gynecology and Obstetrics, Atlanta, GA, US.; Boston University, Dean’s Office, Boston, MA, US.; New York University, Department of Psychiatry, New York, NY, US.; United States Army, Command, Fort Sill, OK, US.; University of Adelaide, Department of Psychiatry, Adelaide, South Australia, AU.; VA Boston Health Care System, GRECC/TRACTS, Boston, MA, US.; Harvard University, Department of Psychology, Boston, MA, US.; UNC Institute for Trauma Recovery, Department of Emergency Medicine, Chapel Hill, NC, US.; Gallipoli Medical Research Institute, PTSD Initiative, Greenslopes, Queensland, AU.; Queensland University of Technology, Faculty of Health, Brisbane, QLD, AU.; Aarhus University Hospital, Psychosis Research Unit, Risskov, DK.; Aarhus University, Centre for Integrated Register-based Research, Aarhus, DK.; Aarhus University, National Centre for Register-Based Research, Aarhus, DK.; University of Copenhagen, Mental Health Services in the Capital Region of Denmark, Mental Health Center Copenhagen, Copenhagen, DK.; National Center for Post Traumatic Stress Disorder, Executive Division, White River Junction, VT, US.; Veterans Affairs San Diego Healthcare System, Department of Research and Psychiatry, San Diego, CA, US.; Northern Illinois University, Department of Psychology, DeKalb, IL, US.; University of California San Diego, Department of Psychiatry, La Jolla, California, US.; University of Texas San Antonio, Department of Psychology, San Antonio, TX, US.; University of Washington, Department of Psychology, Seattle, WA, US.; U.S, Department of Veterans Affairs National Center for Posttraumatic Stress Disorder, West Haven, CT, US.; Yale University School of Medicine, Department of Psychiatry, New Haven, CT, US.; Minneapolis VA Health Care System, Department of Mental Health, Minneapolis, MN, US.; Minneapolis VA Health Care System, Department of Psychology, Minneapolis, MN, US.; University of Minnesota System, Department of Psychiatry, Minneapolis, MN, US.; Charité - Universitätsmedizin, Department of Psychiatry and Psychotherapy, Berlin, GE.; Harvard T.H, Chan School of Public Health, Department of Environmental Health, Boston, MA, US.; Medical University of South Carolina, Department of Nursing and Department of Psychiatry, Charleston, SC, US.; Louisiana State University Health Sciences Center, School of Medicine and Department of Physiology, New Orleans, LA, US.; Maastricht Universitair Medisch Centrum, School for Mental Health and Neuroscience, Department of Psychiatry and Neuropsychology, Maastricht, Limburg, NL.; Universidad Peruana de Ciencias Aplicadas Facultad de Ciencias de la Salud, Department of Medicine, Lima, Lima, PE.; Harvard Medical School, Department of Pediatrics, Boston, MA, US.; University of Michigan, School of Nursing, Ann Arbor, Michigan, US.; University of New South Wales, Department of Psychiatry, Sydney, NSW, AU.; Massachusetts General Hospital, Boston, MA, US.; University of Cape Town, SA MRC Unit on Risk & Resilience in Mental Disorders, Department of Psychiatry, Cape Town, ZA.; Columbia University Medical Center, Department of Medicine, New York, NY, US.; Mental Health Centre Sct, Hans, Institute of Biological Psychiatry, Roskilde, DK.; Oslo University Hospital, KG Jebsen Centre for Psychosis Research, Norway Division of Mental Health and Addiction, Oslo, NO.; University of Illinois at Urbana-Champaign, Carl R, Woese Institute for Genomic Biology, Urbana, IL, US.; University of Illinois at Urbana-Champaign, Department of Psychology, Urbana, IL, US.; Uniformed Services University, Department of Psychiatry, Bethesda, Maryland, US.; Arq, Psychotrauma Reseach Expert Group, Diemen, NH, NL.; Leiden University Medical Center, Department of Psychiatry, Leiden, ZH, NL.; Netherlands Defense Department, Research Center, Utrecht, UT, NL.; New York University School of Medicine, Department of Psychiatry, New York, NY, US.; Amsterdam UMC (location VUmc), Department of Anatomy and Neurosciences, Amsterdam Holland, NL.; Amsterdam UMC (location VUmc), Department of Psychiatry, Amsterdam, Holland, NL.; Medical University of South Carolina, Department of Psychiatry and Behavioral Sciences, Charleston, SC, US.; Ralph H Johnson VA Medical Center, Department of Mental Health, Charleston, SC, US.; University of Copenhagen, Department of Clinical Medicine, Copenhagen, DK.; Durham VA Medical Center, Mental Health Service Line, Durham, NC, US.; James J Peters VA Medical Center, Department of Mental Health, Bronx, NY, US; Baylor Scott and White Central Texas, Department of Psychiatry, Temple, TX, US.; CTVHCS, Psychiatry Research, Temple, TX, US.; Queensland University of Technology, School of Psychology and Counseling, Brisbane, QLD, AU.; Yale University, Department of Biostatistics, New Haven, CT, US.; University of Washington, Department of Psychiatry and Behavioral Sciences, Seattle, WA, US.; Harvard School of Public Health, Department of Epidemiology, Boston, MA, US.

## Abstract

Post-traumatic stress disorder (PTSD) is a common and debilitating disorder. The risk of PTSD following trauma is heritable, but robust common variants have yet to be identified by genome-wide association studies (GWAS). We have collected a multi-ethnic cohort including over 30,000 PTSD cases and 170,000 controls. We first demonstrate significant genetic correlations across 60 PTSD cohorts to evaluate the comparability of these phenotypically heterogeneous studies. In this largest GWAS meta-analysis of PTSD to date we identify a total of 6 genome-wide significant loci, 4 in European and 2 in African-ancestry analyses. Follow-up analyses incorporated local ancestry and sex-specific effects, and functional studies. Along with other novel genes, a non-coding RNA (ncRNA) and a Parkinson’s Disease gene, *PARK2*, were associated with PTSD. Consistent with previous reports, SNP-based heritability estimates for PTSD range between 10-20%. Despite a significant shared liability between PTSD and major depressive disorder, we show evidence that some of our loci may be specific to PTSD. These results demonstrate the role of genetic variation contributing to the biology of differential risk for PTSD and the necessity of expanding GWAS beyond European ancestry.

## Introduction

Posttraumatic stress disorder (PTSD) is a commonly occurring mental health consequence of exposure to extreme, life threatening stress and/or serious injury/harm. PTSD is frequently associated with the occurrence of comorbid mental disorders such as major depression^1^ and other adverse health sequelae including type 2 diabetes and cardiovascular disease.^2,3^ Given this high prevalence and impact, PTSD is a serious public health problem. An understanding of the biological mechanisms of risk for PTSD is therefore an important goal of research ultimately aimed at its prevention and mitigation.^4,5^

Exposure to traumatic stress is, by definition, requisite for the development of PTSD, but individual susceptibility to PTSD (conditioned on trauma exposure) varies widely. Twin studies over the past two decades provide persuasive evidence for at least some genetic influence on PTSD risk,^6,7^ and the last decade has witnessed the beginnings of a concerted effort to detect specific genetic susceptibility variants for PTSD.^8–12^

The Psychiatric Genomics Consortium – PTSD Group (PGC-PTSD) recently published results from the largest GWAS on PTSD to date, involving a transethnic sample of over 20,000 individuals, approximately 25% of whom were cases.^13^ With this limited sample size, no individual variants exceeded genome-wide significance, however, significant estimates of SNP heritability and correlations between PTSD and other mental disorders such as schizophrenia were demonstrated for the first time.

It is apparent from previous PGC work on other mental disorders that sample size is paramount for GWAS to discern common genome-wide significant variants of small effect that are replicable.^14^ Since the publication of data from the first freeze,^13^ the PGC-PTSD has continued to acquire additional PTSD cases and controls through partnerships with an expanding network of investigators, such that we now have accrued a sample size that has enabled us to “turn the corner” on genome-wide risk discovery. Presented here are the results of our latest GWA studies that include over 23,000 European and over 4,000 African ancestry PTSD cases, now involving a total trans-ethnic sample of over 200,000 individuals. In achieving this sample size, both PTSD-associated common variants of small effect in distinct ancestral populations and key genomic pathways associated with risk for PTSD have been identified.

## Results

We report meta-analyses of GWAS from the PGC-PTSD Freeze 2 dataset (PGC2), comprised of an ancestrally diverse group of 206,655 participants. First, a significant genetic signal for PTSD (i.e., genetic correlations) between subsets of studies is demonstrated. Meta-analyses of GWAS including 174,659 individuals (23,212 cases and 151,447 controls) of European ancestry (EUA) and meta-analyses of 15,339 individuals (4,363 cases and 10,976 controls) of African ancestry (AFA) identify a total of 6 independent loci that are significantly associated with PTSD in GWAS of both sexes and stratified by sex. We deeply explore the AFA hits through local ancestry analyses and functional follow-up. Next, we compare genetic heritability of PTSD between sex and ancestries, and show significant genetic risk score predictions for PTSD in different target samples. Finally, we show shared genetic liability of PTSD with related disorders and traits, but confirm that our GWAS hits are specific to PTSD. A detailed description of these results is available in the **Supplementary Note**.

### Meta-analysis strategy across ancestries and sex

Data from 60 different multi-ethnic PTSD studies were included in PGC2 (for study description and demographics see **Supplementary Table 1 and Methods**). Primary GWAS were performed separately for the 3 largest, genetically defined (**Supplementary Figure 1**) ancestry groups (EUA, AFA, Native American Ancestry (AMA), see **Methods**) using logistic regression analyses under an additive genetic model, and meta-analyzed across studies and ancestry groups. Given the previously observed differences between male and female heritability estimates in PGC-PTSD Freeze 1,^13^ we also performed sex-stratified analyses. Quantile-quantile plots showed low inflation across analyses (**Supplementary Figure 4**), which was mostly accounted for by polygenic SNP effects with little indication of residual population stratification (**Supplementary Note**).

### Comparability of PGC2 studies

PGC2 compiled the largest collection of global PTSD GWAS to date, with subjects recruited from both clinically deeply characterized, small patient groups and large cohorts with self-reported PTSD symptoms. We did not restrict the type of trauma subjects were exposed to, and trauma included both civilian and/or military events, often with pre-existing exposure to childhood trauma. To evaluate the comparability of these phenotypically heterogeneous studies we first estimated genetic correlations with LDSC,^15^ a method that leverages GWAS summary results, the only data type available to PGC-PTSD for several of the larger military and non-US cohorts. We found significant genetic correlations (*r_g_*) between studies using a cross-validation approach including all PGC2 EUA subjects (10 runs with studies randomly placed into 2 groups; mean *r_g_* = 0.56, mean SE = 0.23, mean p = 0.029, **Supplementary Table 8**).

Next, additional analyses on the UK Biobank cohort (UKBB) were performed. This cohort comprises a very large proportion of the data, with almost as many EUA cases as the rest of the EUA PGC2 combined (referred to as PGC1.5). PTSD screening in UKBB was based on self-reported symptoms from a mental health survey.^16^ We found a considerable genetic correlation between the UKBB and PGC1.5 EUA subjects (*r_g_* = 0.73, SE = 0.21, p = 0.0005; **Supplementary Table 9**). Further, sensitivity analyses in the UKBB using 3 alternative inclusion criteria for PTSD cases and controls showed stable correlations with PGC1.5 (P1 – P3; *r_g_* = 0.72 - 0.79; **Supplementary Table 10**). Subsequent analyses were based on the UKBB phenotype including the largest number of subjects (P1; N = 126,188). Sex-stratified genetic correlations support the findings of a significant genetic signal across PTSD studies in women and UKBB men, but less so in men from PGC1.5 (discussed in the **Supplementary Note)**.

### Meta-analysis of GWAS in subjects of European and African ancestry

Our largest PTSD meta-analysis in subjects of EUA (N = 174,659) identified two independent, genome-wide significant loci, both mapping to chromosome 6, and sex-stratified analyses in men identified two additional loci (Figures 1A and **1B**, respectively). The smaller meta-analyses in AFA (N = 15,339) identified one genome-wide significant locus, and an additional locus was found in men when stratified by sex (Figures 1C and **1D**, respectively). No genome-wide significant associations were found in meta-analyses of EUA or AFA women (**Supplementary Figure 5**).

**Figure 1.**
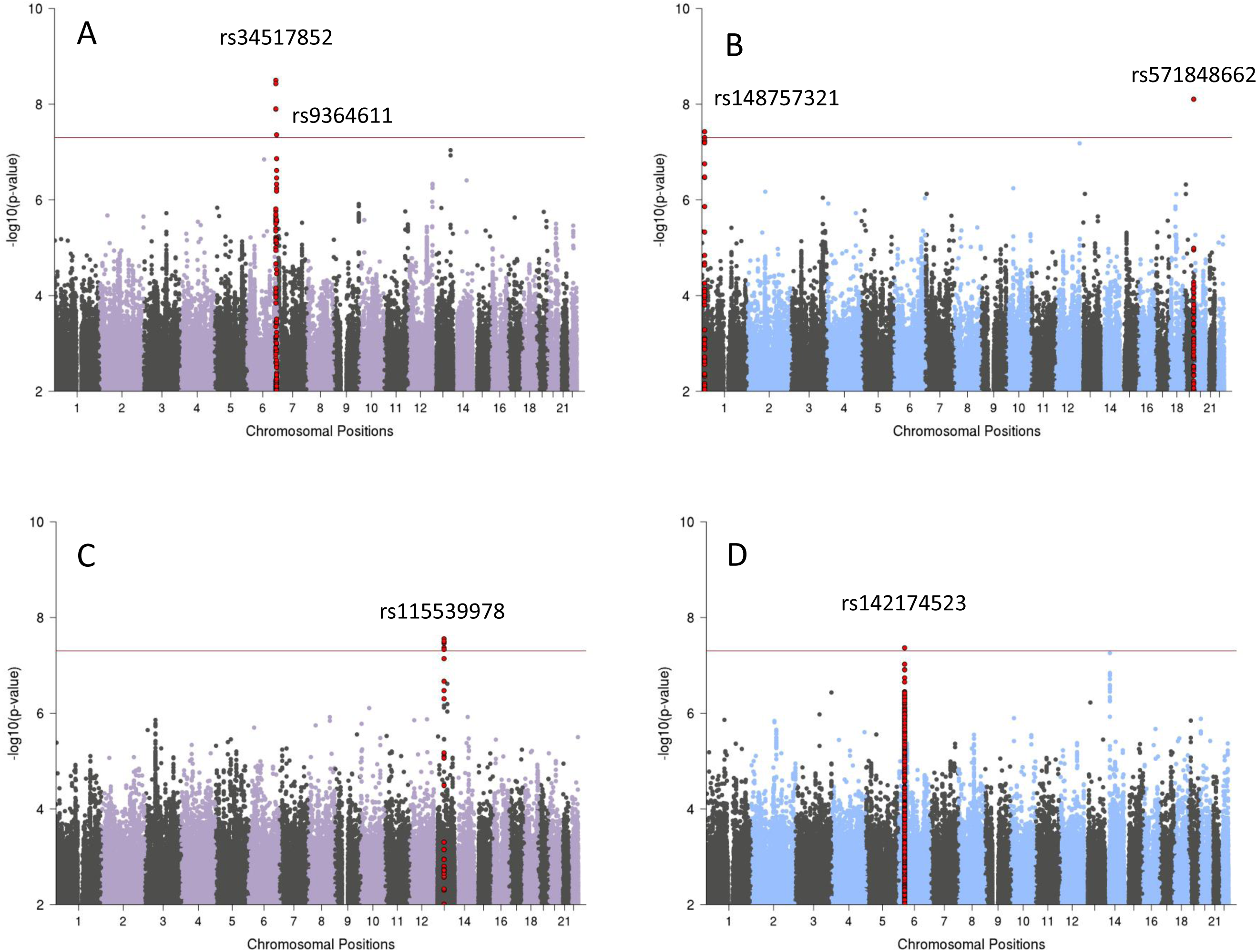
Manhattan plots from meta-analyses of GWAS for PTSD, showing the top variants in 6 independent genome-wide significant loci. Results are shown for subjects of European (EUA; panel A) and African ancestry (AFA; panel C), and for sex-stratified analyses in EUA men (panel B) and AFA men (panel D), respectively. Sex-stratified analyses for women were not significant (Supplementary Fig. 9). The red line represents genome-wide significance at p <5×10^−8^.

Regional plots of the 6 genome-wide significant genomic regions can be found in **Supplementary Figures 6-11.** The 6 leading markers show odds ratios of 1.12 – 1.33 and no significant heterogeneity across studies (**Table 1**). This is supported by PM-plots showing a high consistency of effects among the studies predicted to have an effect^17^ (**Supplementary Figures 12-17**). A z-test on the effect sizes confirmed similar effects for men and women for the 3 leading variants in the joint-sex analyses, and significant sex-specificity for the 3 male hits identified in the sex-stratified analyses (**Supplementary Table 11**).

### Analyses across ancestry groups

In order to study whether the genetic associations with PTSD vary across different ancestries, we first tried to replicate our 6 top hits in the AFA and AMA ancestry groups. No evidence of replication was found by directly comparing the 6 leading markers, nor by investigating the larger genomic regions harboring the signal (**Supplementary Figures 18-23**). In addition, we did not identify any genome-wide significant hits by performing a trans-ethnic genome-wide meta-analysis across the 3 main ancestry groups under fixed-and random-effect models (**Supplementary Figure 24**).

While lack of replication of the 4 EUA hits may not be conclusive due to limited power in the smaller AFA and AMA data (**Supplementary Tables 4-5**), the 2 hits in AFA provided an opportunity for a more detailed analysis of ancestry effects. GWAS in the AFA subjects included standard PCs to control for average admixture across the genome. However, to precisely infer local ancestry across the genome of admixed subjects we further implemented a local ancestry inference (LAI) pipeline (**Supplementary note**). We confirmed the AFA top hit rs115539978 to be specific to the African ancestral background (8% MAF on the African, and <1% MAF on the European and Native American backgrounds, respectively; **Supplementary Table 12**). Conversely, LAI analyses of the male-specific hit indexed by rs142174523 showed no evidence for ancestry-specific effects that would explain the lack of replication in the EUA meta-analyses; however, the LD-structure in the MHC locus is complex.^18^

### Integration with functional genomic data

Functional mapping and annotation of the 6 GWAS hits using the FUMA pipeline conservatively predicted 5 genes *ZDHHC14, PARK2, KAZN, TMRM51-AS1,* and *ZNF813* located in EUA risk loci, and 5 distinct genes *LINC02335, MIR5007, TUC338, LINC02571,* and *HLA-B* in AFA risk loci (**Table 2**). Gene-based analyses on 18,222 protein-coding genes identified 2 additional gene-wide significant loci, represented by *SH3RF3* (p = 4.28−10^−07^) and *PODXL* (p = 2.37×10^−06^) in the combined EUA analysis. Many of these new potential target genes are very interesting with respect to our understanding of the biology of PTSD. The biological function and potential psychiatric relevance of these 12 genes is detailed in the **Supplementary Note**.

We next performed gene-set analyses to understand implicated genes in the context of pathways and found 4 significant, Bonferroni corrected gene sets (**Supplementary Table 13).** Of note, the two gene sets identified in the EUA data point towards a role for the immune system in PTSD. This is supported by a number of TNF-related genes summarized in a significant gene set in AFA.

Annotation of variants in risk loci showed limited evidence of functionality (**Table 2** and **Supplementary Note).** Most notably for the AFA top hit on chromosome 13, when testing for chromatin interactions using Hi-C data in neural progenitor cells, significant chromatin conformation interactions were observed between the risk region and a region approximately 1,100kb upstream harboring additional non-coding RNAs including LINC00458, hsa-mir-1297 and LINC00558 as well as a region approximately 820kb downstream harboring the pseudogene *HNF4GP1* (**Supplementary Figure 26**). eQTL analyses did not show significant associations with gene expression. However, the lack of functional data for this region may be explained by the African ancestry specificity of the GWAS findings since databases available within the FUMA framework, including GTEx and BrainSpan for eQTL analyses, are predominantly based on European populations.

Thus, we expanded our analyses for the AFA top hit through cell culture experiments in lymphoblastoid cell lines (LCLs) from African subjects (see **Methods** and **Supplementary Note**). We show evidence that the African-ancestry specific SNP rs115539978 seems to capture a genomic region that may influence the expression of non-coding RNAs from this PTSD risk locus in response to increased glucocorticoid receptor signaling, thus linking this African-specific genetic variant to stress response and non-coding RNA expression (**Supplementary Figure 27**).

We further characterized the AFA signal (rs115539978) using psychophysiology and imaging datasets available through the Grady Trauma Project (GTPC) and found evidence that this lead SNP captures a genomic region that is also associated with increased amygdala volume and fear psychophysiology in a traumatized population (**Supplementary Note and Supplementary Figure 28**).

### Heritability of PTSD

We estimated heritability of PTSD based on information captured by genotyping arrays (*h^2^_snp_*) from summary statistics using LDSC in the EUA studies (**Table 3A**). Assuming a population prevalence of 30% after trauma exposure, *h^2^_snp_* was 0.05 on the liability scale (p = 3.18 × 10^−8^). Female heritability was highly significant (*h^2^_snp_* = 0.10, p = 8.03 × 10^−11^), while male heritability was not significantly different from zero (*h^2^_snp_* = 0.01, p = 0.63).

We further examined these sex-based differences separately in PGC1.5 and UKBB. Again, heritability in PGC1.5 was high in women and not significant in men. In contrast, in the UKBB, male heritability was significant (*h^2^_snp_* = 0.15, p = 1.38 × 10^−3^) and not significantly different (p = 0.41) from heritability in UKBB women (*h^2^_snp_* = 0.19, p = 2 × 10 ^−10^). Heritability estimates using different case and control definitions in UKBB further confirm these results (**Supplementary Table 14**).

We also compared heritability across ancestries using the GCTA GREML method, which allows analysis of admixed populations when individual genotypes are available. GCTA estimates in the smaller EUA data remained similar to LDSC on the full data (**Table 3B**). Heritability in AFA was comparable to estimates for EUA in the overall sample and when stratified by sex.

In summary, with the notable exception of PGC1.5 men, which fail to show significant heritability estimates, overall *h^2^_snp_* for PTSD is in the range of 10-20%, and estimates remain relatively stable across methods, studies, and ancestries.

### Polygenic risk scores (PRS) for PTSD

Genetic risk scores generated from well-powered GWAS have recently become a tool of high relevance for polygenic disorders and traits (e.g. ^19,20^). We assessed the predictive value of the PRS for PTSD developed from our largest EUA sample, the UKBB, and showed a significant increase in odds to develop PTSD across PRS quintiles in PGC1.5 EUA target samples, with a variance explained on the liability scale of *r^2^* = 0.0015 (p = 5.44 × 10^−7^) (Figure 2). The highest OR was seen in UKBB men with a PRS trained on women from the same study, reaching an OR of 1.39 in the 5^th^ quintile and a variance explained of *r^2^* = 0.012 (p = 4.19 × 10^−10^).

**Figure 2.**
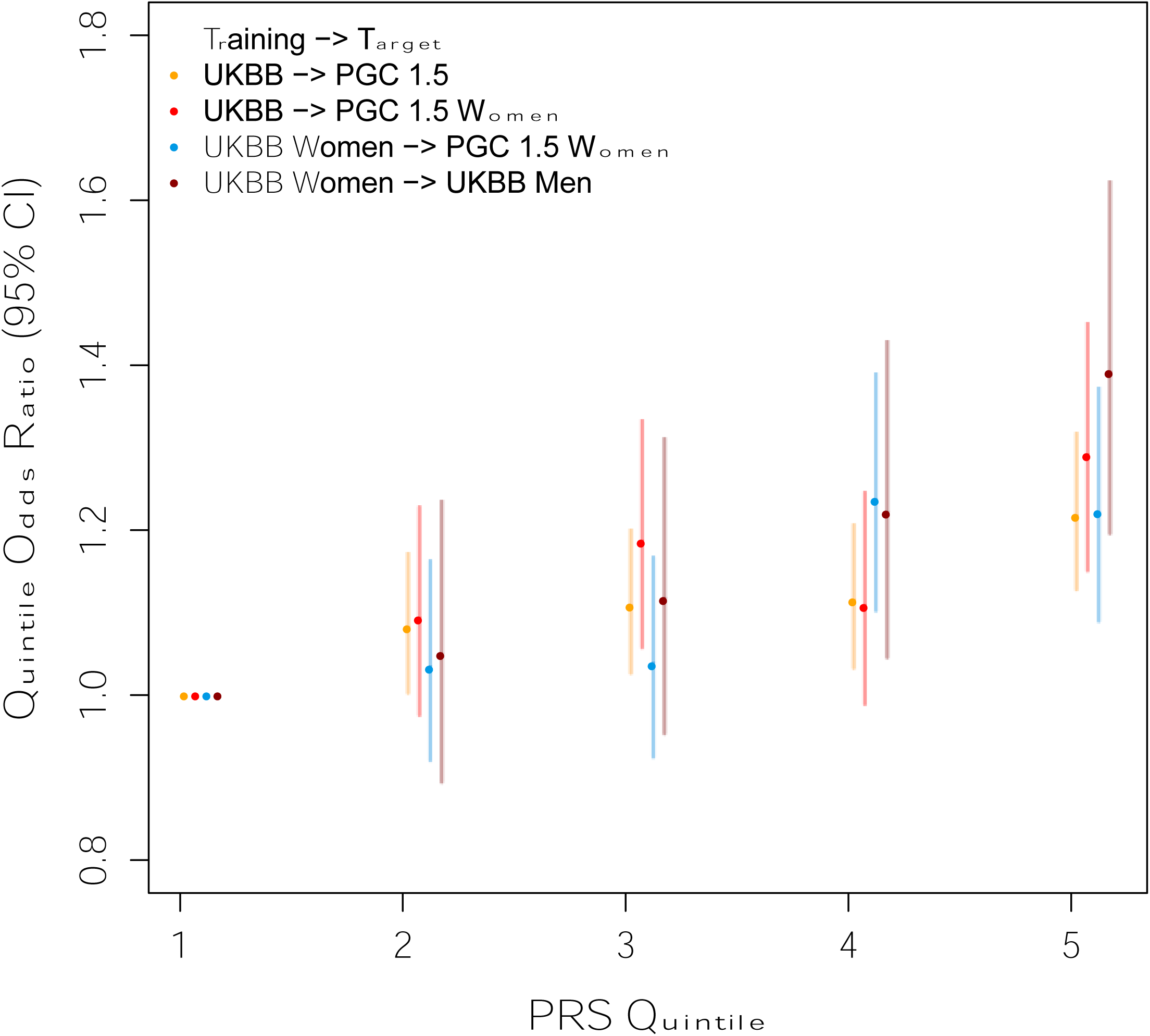
Genetic risk score (PRS) predictions PTSD. Using PTSD subjects from the UK Biobank as discovery sample, odds ratios (OR) for PTSD per PRS quintile relative to the first quintile show a significant increase in different PGC PTSD target samples. For example, UK Biobank men in the 5^th^ quintile have 40% higher odds to develop PTSD than UK Biobank men in the lowest quintile, when using women from the same population as a training set. Sample sizes in different training and target sets: UKBB: 10,389 PTSD, 115,799 controls; UKBB women: 6,845 PTSD, 64,099 controls; UKBB men: 3,544 PTSD, 51,700 controls; PGC1.5: 10,213 PTSD, 27,445 controls; PGC 1.5 women: 5,232 PTSD, 6,551 controls.

### Genetic correlations of PTSD with other traits and disorders

Analysis of shared heritability across common disorders of the brain^21^ and specific genetic correlations of psychiatric disorders with cognitive, anthropomorphic and behavioral measures^13,22–24^ has been facilitated greatly by the development of a centralized database of GWAS results including a web interface for LD score regression (LD Hub^25^). We estimated pairwise genetic correlations (*r_g_*) between PTSD and 235 disorders/traits and found 21 significant correlations after conservative Bonferroni correction (Figure 3, **panel A and Supplementary Table 15**). Genetic variation associated with PTSD was positively correlated with PRS from other psychiatric traits including depressive symptoms, schizophrenia and neuroticism, as well as epidemiologically comorbid traits such as insomnia, smoking behavior, asthma, hip-waist ratio and coronary artery disease. In contrast, negative *r_g_* with PTSD include subjective well-being, education, and a strong correlation with parents age at death (*r_g_* = -*0.70)*. Significant positive correlations were also found for reproductive traits such as the number of children ever born, and, as previously reported for women,^26^ PTSD was negatively correlated with age at first birth.

**Figure 3.**
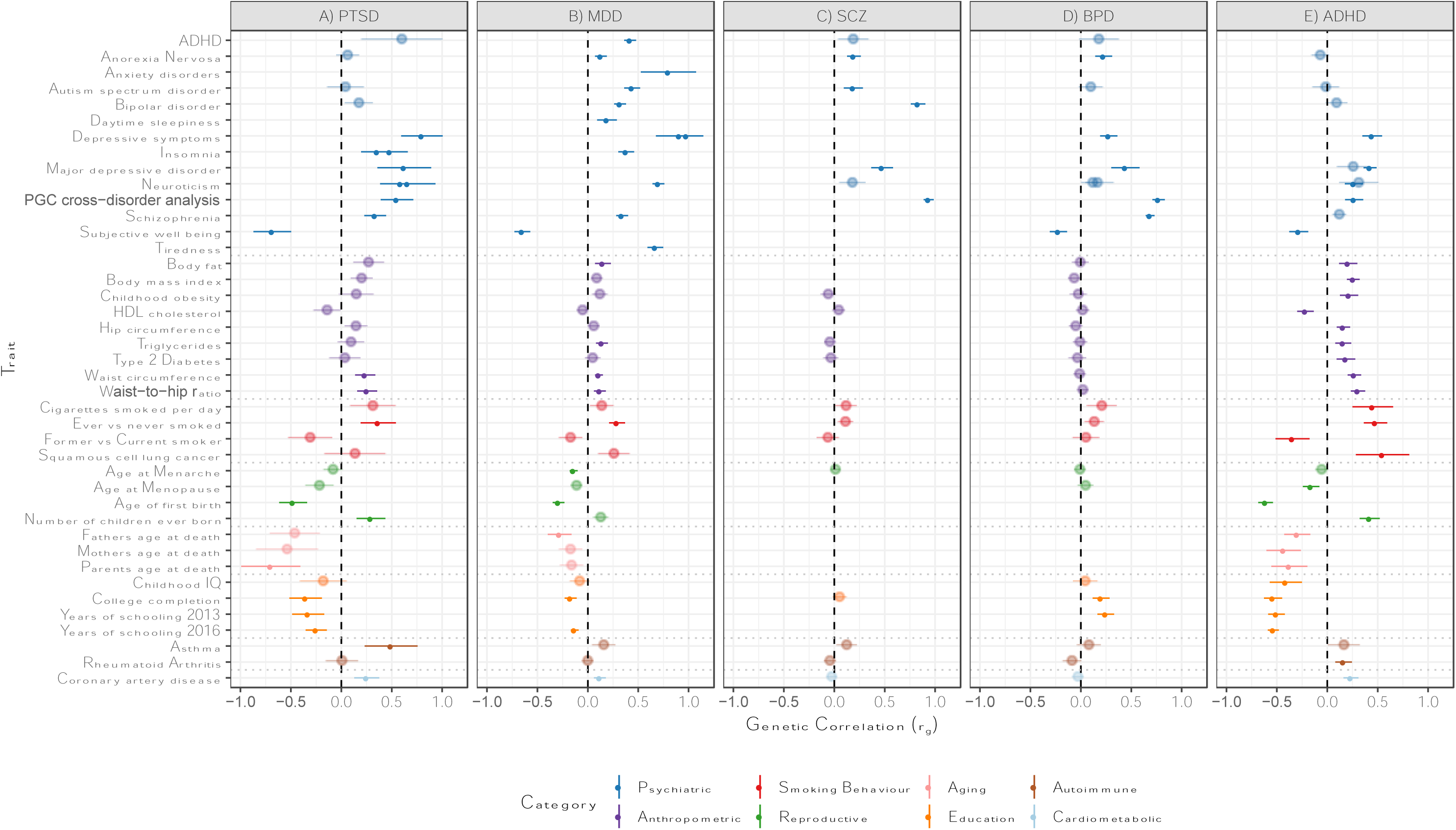
Genetic correlations between several groups of traits (from psychiatric, anthropomorphic, smoking behavior, reproductive, aging, education, autoimmune, and cardiometabolic categories) and A) PTSD, B) MDD, C) SCZ, D) BPD, and E) ADHD. Only traits with at least one significant correlation with the 5 psychiatric disorders are shown. Error bars indicate 95% confidence limits. Solid points indicate significant correlation after Bonferroni correction. The total number of correlations tested were 235 for PTSD, 221 for MDD, 172 for SCZ, 196 for BPD, and 219 for ADHD. Due to substantial overlap with other phenotype traits, 2 education, 4 anthropometric, and 2 cancer phenotypes were omitted.

With the notable exception of asthma, our findings on PTSD correspond closely with genetic correlations between these traits and other psychiatric disorders such as MDD^22^, SCZ^13^, BPD^23^, and ADHD^24^ (Figure 3, **panels B-E**). These findings are not surprising, as pleiotropic effects (i.e. SNPs impacting multiple traits) have been widely reported for psychiatric disorders.

In order to test if the genome-wide significant findings from our GWAS meta-analyses were specific to PTSD, we adjusted the top hits from our analyses for the effects of genetically correlated psychiatric traits. Since the strongest correlations were found between PTSD and depressive symptoms *(*r_g_* = 0.80)* and MDD *(r_g_ = 0.62)*, summary statistics from PGC MDD ^22^, as well as MDD plus BPD and SCZ, were included in the analyses. Using a recently implemented method for GWAS summary data (mtCOJO^27^), we found that effect sizes for the 4 EUA top hits were not markedly reduced when adjusted for the effects of MDD, or all 3 psychiatric traits tested (**Supplementary Table 16a and b**). These findings indicate that the genetic variants identified here are specific to PTSD when tested in the context of the 3 genetic disorders most significantly correlated with PTSD.

## Discussion

PTSD is a common and debilitating condition known to be influenced by genetic factors, yet few common genetic risk variants for PTSD have been robustly identified. The PGC-PTSD combined data from 60 multi-ethnic cohorts (PGC1.5) and the UK Biobank to produce the largest effort to date (PGC2), a sample size of 206,655 participants, over ten times that of any previous analysis.^13,28^ As has been demonstrated in GWAS of SCZ,^29^ BPD,^30^ and recently in MDD^31^ and ADHD,^24^ sample size is critical to produce robust genome-wide significant hits that can inform foundational knowledge on the neurobiology of complex psychiatric conditions. These results show this is also true of PTSD. This increased power has led us to draw several major conclusions.

First, our genetic findings squarely place PTSD among the other psychiatric disorders. While this statement may seem obvious to some, there remains debate about whether PTSD is entirely a social construction.^32^ We found substantial SNP-based heritability (i.e., phenotypic variation explained by genetic differences) at 10-20%, similar to that for major depression^22^ across methods, studies and ancestries. The heritability results and pattern of genetic correlations are also consistent with our initial findings^13^ and with those from twin studies. PTSD shares common variant risk with other psychiatric disorders, which show substantial sharing of common variant risk with one another.^33^ PTSD was most significantly (genetically) correlated with major depression, but also with schizophrenia, both of which have genome-wide significant loci implicated in brain function.

Second, our ancestry specific analyses identified several novel genetic loci associated with PTSD. Two independent genome-wide significant loci, both mapping to chromosome 6, were found in our largest meta-analysis of EUA subjects and two additional loci were identified in sex-stratified analyses in EUA men only. A smaller meta-analysis in AFA identified one genome-wide significant locus, which we determined to be ancestry-specific specific though local ancestry analyses, and an additional locus was found in AFA men when stratified by sex. These loci pointed to a number of different target genes that are very interesting with relationship to our understanding of the biology of PTSD. With *PARK2,* there is a posited role of dopaminergic systems in PTSD and epidemiological evidence for association of Parkinson Disease and PTSD.^34–36^ *PODXL* is involved in neural development and synapse formation,^37,38^ *SH3RF3* is associated with neurocognition and dementia,^39,40^ and *ZDHHC14* is associated with regulation of (B-adrenergic receptors.^41,42^ Finally, the *HLA-B* complex may be related through the known role of immunity and inflammation in stress-related disorders.^43,44^ Extensive follow-up work is needed to determine the function of identified genes and their relationship to putative pathological processes. For example, in SCZ the MHC locus is now thought to influence risk through pruning of synapses using immune machinery rather than through classical immune pathways.^45^ These ancestry-specific results are preliminary, and even larger PTSD GWAS will facilitate the identification of plausible neurobiological targets for PTSD.

Third, our results also illustrate that there may be genetic contributions to the well-documented association between PTSD and dysregulation in inflammatory and immune processes.^46^ It has been widely recognized that PTSD is associated with a broad range of adverse physical health conditions over the life course ranging from type-2-diabetes and cardiovascular disease to dementia and rheumatoid arthritis.^47,48^ Less is known about the biological mechanisms driving the relationship between PTSD and these outcomes. Our genetic correlation analyses may provide some initial clues for further investigation. For example, we found a high genetic correlation (*r_g_* = 0.49, p = 0.0002) between PTSD and asthma. Our subsequent gene set and pathway analyses provide some clues further implicating the immune system. Of note, these genetic results converge with evidence from epidemiologic cohort studies documenting the role of stress-related disorders such as PTSD in autoimmune diseases,^49^ case-control studies showing elevations of immune-related biomarkers in women with PTSD,^50,51^ and epigenetic studies pointing to the role of the immune system in PTSD etiology.^52–54^ Of note, it has been previously suggested that PTSD may be “a systemic disorder rather than one confined to the mind”.^55^ Further work is needed to determine whether PTSD has genetic overlap with immune disorders broadly and the causal direction between disorders. At minimum, the emerging genetic evidence presented here suggests that association between PTSD and health conditions may, in some cases, have some genetic origin.

Fourth, PGC-PTSD is distinct in relation to current genomics consortia due to its high proportion of data from participants of diverse ancestries. For example, a recent review found that only three percent of all samples in genetic studies were from African ancestry.^56^ This contrasts sharply with the 10% of AFA participants in our consortium. We have the first heritability estimates for PTSD in African ancestry: they are similar to EUA, highly significant in women and lower in men. Our GWAS in subjects of African ancestry indicated at least one ancestry-specific locus using local ancestry methods developed for this analysis. We note the sample size in the AFA analysis has only about 15,000 participants, which is small and under-powered. This finding could be a false positive. However, other work has shown that genetic studies of underrepresented populations afford the opportunity to discover novel loci that are invariant in European populations.^57^ As others have noted, there are major limitations in our knowledge of the genetic and environmental risk architecture of psychiatric disorders in persons of African descent.^58^ Our findings provide further evidence of the need to invest in research that includes diverse ancestral populations, to expand reference data, and to continue to develop methods to analyze data from such populations. Until such an investment is made, we are limited in our ability to understand biological mechanisms, predict genetic risk,^59^ and produce optimal therapy for non-European populations. African genomes are characterized by shorter haplotype blocks and contain many millions more variants per individual than populations outside Africa.^60^ Further, including data from African populations in genetic studies of PTSD and other neuropsychiatric disorders may accelerate genetic discovery and could be useful for fine mapping disease causing alleles.^61^

Fifth, although PTSD heritability remained relatively stable across methods, studies, and ancestries, sex differences in heritability were observed in the overall cohort analyses as well as in the AFA analyses.^13^ It is of note that the sex differences in heritability were not evident in the UK Biobank data, which we hypothesize is due to two differences between the PGC1.5 cohorts and the UK Biobank. PTSD is conditional on trauma exposure, which is also highly heterogeneous across individuals and populations.^62^ Unlike PGC1.5, the UK Biobank cohort is comprised of few to no subjects who experienced military-related trauma. In contrast, a substantial proportion of men in the PGC1.5 cohorts were from military cohorts, while virtually all women were civilians. The environmental experiences (e.g., military versus civilian) and index traumatic events leading to PTSD in male subjects versus female subjects could explain observed lower heritability estimates in males in the PGC1.5 cohorts. In future work, we aim to investigate this empirically by pooling detailed trauma and PTSD phenotypic information on both males and females and by modeling the effects of measurement variability on heritability estimates. Future work could also aim to increase samples of civilian men and military women to allow for analyses stratified by military trauma and sex.

Finally, we now have a significant polygenic risk score for PTSD. Larger sample sizes are needed to increase the variance explained and improve sensitivity and specificity.^20^ In the future, polygenic risk may eventually be useful in algorithms developed to identify vulnerable persons after exposure to trauma. PTSD is one of the most preventable mental disorders, as many people exposed to trauma come to clinical attention in first response settings such as emergency rooms, intensive care units, and trauma centers. Controlled clinical trials show that PTSD risk can be significantly reduced by early preventive interventions.^63–65^ However, these interventions have nontrivial costs, making it infeasible to offer them to all persons exposed to trauma, given that only a small minority goes on to develop PTSD.^66–69^ They are also unnecessary for many survivors who recover spontaneously.^64^ To be cost-effective, risk prediction rules are needed to identify which exposed persons are at high risk of PTSD. Such risk prediction tools have been developed,^70,71^ but none to date has included polygenic risk as a predictor.

These findings substantively advance our understanding of the genetic basis of PTSD, and they also demonstate that PGC-PTSD remains under-powered for the detection of most risk loci. PTSD is similar to major depression in both prevalence (among trauma-exposed persons) and in heritability. There are now 44 genome-wide significant hits for major depression; that level of discovery required over 135,000 cases and 344,000 controls.^31^ Other limitations include the treatment of PTSD as a binary disorder in our analysis. Extensive epidemiologic work has shown that subthreshold PTSD is highly prevalent and debilitating.^72,73^ In our analysis, persons with subthreshold PTSD are classified as controls, which would likely reduce our power to find genetic associations. In future work, we will consider PTSD as a continuous phenotype as well as examine clusters of PTSD symptoms, which are more homogeneous; Gelernter et al. (2018) found multiple genome-wide significant loci for re-experiencing symptoms, which is the cluster of symptoms most unique to PTSD, in data from over 100,000 veterans in the Million Veterans Program.^74^ Finally, we used mostly unscreened controls, but controls carefully screened for trauma may increase power since trauma is required for a PTSD diagnosis. In addition to increasing sample size, we aim in the future to also pool item-level phenotypic data from our cohorts in order to address these limitations.

## Online Methods

### Participating studies

The PGC-PTSD Freeze 2 dataset (PGC2) included 60 ancestrally diverse studies from Europe, Africa and the Americas. Of these, 12 were already included in Freeze 1.^13^ Study details and demographics can be found in **Supplementary Table 1**. PTSD assessment was based either on lifetime (where possible) or current PTSD (i.e., including participants with a potential lifetime PTSD diagnosis as controls), and PTSD diagnosis was established using various instruments and different versions of the DSM (DSM-III-R, DSM IV, DSM-5). For GWAS analyses, all studies provided PTSD case status as determined using standard criteria and control subjects not meeting the PTSD diagnostic criteria (see **Supplementary Table 1** for additional exclusion criteria).

The following studies were included, listed with the official name of each study followed by a 4-letter abbreviation corresponding to genotyping files and a study number corresponding to **Supplementary Table 1**:

### Army Study to Assess Risk and Resilience in Servicemembers (NSS1, NSS2, PPDS; Supplementary Table 1 #14-16)

See reference for details.^75^ Potentially traumatic events in childhood and adult civilian trauma were assessed in all participants, as well as military traumatic experience for participants who had been deployed, using a self-administered questionnaire. The extent of traumatic experiences was summarized, separately, in a non-deployment trauma variable and a deployment trauma variable. Both continuous variables are summaries of frequencies of responses to each of the 11 questions regarding non-deployment or deployment trauma. Responses range from Never (0), Once (1), 2-4 Times (2), 5-9 Times (3) or More than 10 Times (4). A combination of a computer-administered version of the Composite International Diagnostic Interview (CIDI) and the Posttraumatic Stress Disorder Checklist (PCL for DSM-IV)^76^ were used to assign diagnoses of lifetime Posttraumatic Stress Disorder (PTSD) according to DSM-IV.^77^ DNA for GWAS analysis was isolated from blood. The Institutional Review Boards of all participating institutions approved this study.

### Ash Wednesday (BRYA; Supplementary Table 1 #10)

See reference for details.^78^ Potentially traumatic events were identified using the Recent Life Events questionnaire.^79^ The PTSD Checklist (PCL) was used to assess PTSD over the prior 4 weeks by interviews.^76^ Respondents were considered to have a diagnosis if DSM-IV criteria were met. The PCL calculates PTSD symptom severity, which ranged from 0 to 68, by clinical interview. For this cohort (187 Cases, 261 Controls), the mean severity was 3.28 and the standard deviation 11.56. DNA for GWAS analysis was isolated from saliva. The Institutional Review Board of Western Sydney Area Health Service approved this study.

### Biological Effects of Traumatic Experiences, Treatment and Recovery (BETR; Supplementary Table 1 #48)

See reference for details.^80^ In total, 57 PTSD patients, 29 veteran controls (combat controls) and 32 civilian controls (healthy controls) were included. Patients were recruited from one of four outpatient clinics of the Military Mental Healthcare Organization, The Netherlands. Patients were included after a psychologist or psychiatrist diagnosed PTSD. PTSD diagnosis was confirmed using the Clinician Administered PTSD scale (CAPS ≥45^81^). The Structural Clinical interview for DSM-IV (SCID-I^82^) was applied to diagnose comorbid disorders. Control participants were recruited via advertisements, and the interviews (SCID and CAPS) were also applied to investigate PTSD symptoms and psychiatric disorders. Inclusion criteria for controls were no current psychiatric or neurological disorder, and no presence of current PTSD symptoms (CAPS ≤15). After receiving a complete written and verbal description of the study all participants gave written informed consent. The Medical Ethical Committee of the UMC Utrecht approved the study, and the study was performed in accordance with the Declaration of Helsinki.

Potentially traumatic events were identified using the Life Events Checklist for DSM-IV (LEC-IV).^83^ The CAPS for DSM-IV was used to assess PTSD over the prior month by trained researchers.^81^ The CAPS calculates PTSD symptom severity, which ranged from 0 to 107, by calculating the total CAPS score. For this cohort (57 Cases, 61 Controls), the mean severity was 50.05 and the standard deviation 32.86. Respondents were considered to have a lifetime diagnosis if DSM-IV criteria were met. Respondents were considered to have a current diagnosis if DSM-IV were met in the previous month. The Institutional Review Board of Utrecht University Medical Center approved this study.

### Bounce Back Now (BOBA; Supplementary Table 1 #18)

See reference for details.^84^ Potentially traumatic events were identified using National Survey (NSA) on Adolescents PTSD module which was administered by trained interviewers using computer assisted telephone interview technology. The NSA PTSD Module assessed for exposure to five types of potentially traumatic events, in addition to five specific questions about the impact of the tornado (e.g., did the tornado cause damage to your house or property?).^84,85^ The NSA PTSD Module that was administered was used to assess PTSD since the tornado, as well as during any two week period in their lifetimes, and lifetime PTSD was used in the present analyses.^85^ Respondents were considered to have a lifetime diagnosis if DSM-IV PTSD criteria (i.e., at least one re-experiencing symptom, three avoidance symptoms, and two or more arousal symptoms) were met for at least a two week period. For this cohort, 127 individuals with lifetime PTSD were matched by sex and race/ethnicity with 127 controls who did not meet criteria for lifetime PTSD. DNA for GWAS analysis was isolated from saliva collected via Oragene kits. The Institutional Review Board of the Medical University of South Carolina approved this study.

### Childhood Trauma Study (QIMR; Supplementary Table 1 #30)

See references for details.^86,87^ Potentially traumatic events were identified using a semi-structured psychiatric diagnostic telephone assessment that included questions on childhood maltreatment.^88^ Lifetime DSM-IV PTSD (binary measure) was assessed using a modified version of the measure from the National Comorbidity Survey by telephone interviewers trained by an experienced clinical psychologist.^89^ For respondents who had experienced more than one potentially traumatic event, assessment of lifetime PTSD focused on the event identified by each respondent as most disturbing. DNA for GWAS analysis was isolated from blood. The Queensland Institute of Medical Research Ethics Committee and the Washington University School of Medicine Human Research Protection Office approved this study.

### Child Trauma and Neural Systems Underlying Emotion Regulation (KMCT; Supplementary Table 1 #19)

Potentially traumatic events were identified using The UCLA PTSD Reaction Index,^90^ the Childhood Experiences of Care and Abuse Interview,^91^ and the Childhood Trauma Questionnaire.^92^ The Clinician Administered PTSD Scale for Children^93^ was used to assess both lifetime and current PTSD by trained clinical interviewers.^91^ Children and a parent or guardian completed the interview, and an or rule was used to assign diagnoses. Respondents were considered to have a diagnosis if DSM-5 criteria were met. Respondents were considered to have a current diagnosis if DSM-5 criteria were met. The UCLA PTSD Reaction Index^90^ calculates PTSD symptom severity. Children and a parent or guardian each completed this measure, and we used the highest score from either the child or parent, which ranged from 0 to 67 in our sample. For this cohort (133 Cases, 122 Controls), the mean severity was 17.57 and the standard deviation 17.94. DNA for GWAS analysis was isolated from saliva. The Institutional Review Board of the University of Washington approved this study.

### CHOICE (FEEN; Supplementary Table 1 #37)

See reference for details.^94–103^ Potentially traumatic events were identified using the standard trauma interview.^104^ The PTSD Symptom Scale – Interview (PSS-I) was used to assess PTSD over the prior two weeks for the trauma of interest by postdoctoral or graduate level assessors trained to reliability^82^. The Structured Clinical Interview (SCID-IV) was used to assess lifetime PTSD (not current) for a trauma not the focus of treatment by postdoctoral or graduate level assessors trained to reliability.^105^ Respondents were considered to have a current diagnosis if on the PSS-I they met symptom-level DSM-IV diagnostic criteria. The PSS-I also provides PTSD symptom severity, with a range from 0 to 51. For this cohort (104 Cases), the mean severity was 32.63 and the standard deviation 4.87. DNA for GWAS analysis was isolated from blood. The Institutional Review Board of University Hospitals approved this study.

### Cohen Veterans Center Study (COM1; Supplementary Table 1 #50)

See reference for details.^106–108^ This is a multi-site study that ran through NYUMC and Stanford University and Palo Alto VAMC. The VA Palo Alto Health Care System has an affiliation with Stanford University School of Medicine. Dr. Charles Marmar is the overall PI for this study.

Potentially traumatic events were identified using clinical interview that was administered by a licensed psychologist.^109^ The CAPS was administered by clinicians to assess PTSD for two time periods: the preceding 30 days and a one-month period in the past when symptoms were the worst, by respondent’s subjective account.^81^ The CAPS *calculates PTSD symptom severity by summing the scores for all items, which ranged from 0 to 80 in the full range* and ranged from 0 to 56 in this dataset (in the Cohen Veterans Center study). For this cohort (232 Cases, 802 Controls), the mean severity was 12.14 and the standard deviation 11.85. Respondents were considered to have a current diagnosis if DSM-5 criteria were met in the preceding 30 days. Respondents were considered to have a lifetime diagnosis if DSM-5 criteria were met in the month when symptoms were the worst. The Institutional Review Board of NYUMC and Stanford University approved this study.

### Cortical Excitability: Biomarker and Endophenotype in Combat Related PTSD (WANG; Supplementary Table 1 #59)

Potentially traumatic events were identified using CAPS Life Event Checklist. The CAPS was used to assess PTSD over the prior 4 weeks, by research coordinators/interviewers.^81^ Respondents were considered to have a diagnosis if CAPS>45. Respondents were considered to have a current diagnosis if CAPS>45. The CAPS calculates PTSD symptom severity, which ranged from 0 to 123 out of a possible 136, by interview.^81^ For this cohort (208 Cases, 87 Controls), the mean severity of cases was 76.22 and the standard deviation was 16.85. DNA for GWAS analysis was isolated from whole blood. The Institutional Review Board of Medical University of South Carolina approved this study.

### Danish military study (DAMI; Supplementary Table 1 #28)

Potentially traumatic events were identified with 11 single items listing potentially traumatic events occurring during deployment. A scale developed for the Danish military, the PRIM-PTSD, was used to assess PTSD-symptoms over the previous 3 months^110^. The PRIM-PTSD calculates PTSD symptom severity, with a possible range of 12-48. For this cohort (462 Cases, 2019 Controls after quality control), the mean severity was 17.97 and the standard deviation 5.92. PTSD cases were defined as having a PRIM-PTSD score at or above 25, equaling a score of 44 on the PTSD Checklist. DNA for GWAS analysis was isolated from neonatal blood spots. The Regional Committee on Health Research Ethics, Region Zealand, approved this study.

### Danish iPSYCH PTSD samples (DAIP; Supplementary Table 1 #29)

See reference for details.^111–113^ The Danish iPSYCH PTSD samples were identified for analysis using infrastructure provided by the iPSYCH project.^114^ The iPSYCH project is a case cohort study, drawing individuals born in Denmark between 1981 and 2005, and obtaining all cases diagnosed with six disorders plus 30,000 random individuals from the same population cohort as controls. PTSD was not one of the original six diagnoses within iPSYCH, but cases were identified in linked records (i.e. PTSD diagnosis comorbid with one of the six ascertained disorders or PTSD diagnosis in one of the 30,000 iPSYCH “controls”). PTSD was assessed via clinician diagnosis according to ICD-10 (F43.1), as obtained from either of two registers in Denmark: the Danish Psychiatric Central Research Register and/or the Danish National Patient Register. Diagnoses in the registers are for current disorders (i.e. not lifetime). PTSD severity was not available. DNA for GWAS analysis was isolated from bloodspots from the Danish Neonatal Screening Biobank hosted by the Statens Serum Institut, as described previously.^112,113^ The study was approved by the Regional Danish Ethics Committee and the Danish Data Protection Agency.

### DCS Rothbaum Study (DCSR; Supplementary Table 1 #38)

See reference for details.^115,116^ The authors examined the effectiveness of virtual reality exposure augmented with D-cycloserine or alprazolam, compared with placebo, in reducing PTSD due to military trauma.

After an introductory session, five sessions of virtual reality exposure were augmented with D-cycloserine (50 mg) or alprazolam (0.25 mg) in a double-blind, placebo-controlled randomized clinical trial for 156 Iraq and Afghanistan war veterans with PTSD.^117^

PTSD symptoms significantly improved from pre- to posttreatment across all conditions and were maintained at 3, 6, and 12 months. There were no overall differences in symptoms between D-cycloserine and placebo at any time. Alprazolam and placebo differed significantly on the Clinician-Administered PTSD Scale (CAPS) score at posttreatment and PTSD diagnosis at 3 months posttreatment; the alprazolam group showed a higher rate of PTSD (82.8%) than the placebo group (47.8%).^81^ Between-session extinction learning was a treatment-specific enhancer of outcome for the D-cycloserine group only. At posttreatment, the D-cycloserine group had the lowest cortisol reactivity and smallest startle response during virtual reality scenes.

A six-session virtual reality treatment was associated with reduction in PTSD diagnoses and symptoms in Iraq and Afghanistan veterans, although there was no control condition for the virtual reality exposure. There was no advantage of D-cycloserine for PTSD symptoms in primary analyses. In secondary analyses, alprazolam impaired recovery and D-cycloserine enhanced virtual reality outcome in patients who demonstrated within-session learning. D-cycloserine augmentation reduced cortisol and startle reactivity more than did alprazolam or placebo, findings that are consistent with those in the animal literature.

### Defining Essential Features of Neural Damage (DEFE; Supplementary Table 1 #12)

See reference for details.^118,119^ Potentially traumatic events were identified using the Clinician-Administered PTSD Scale for DSM-IV (CAPS-IV), the Combat Exposure Scale (CES), and items from the Deployment Risk and Resilience Inventory (DRRI).^81,109,120,121^ The CAPS-IV was also used to assess PTSD currently (over the prior 2 weeks) and over the lifetime by trained interviewers.^81^ The CAPS-IV calculates PTSD symptom severity, which ranged from 0 to 120. For this cohort (88 Cases, 62 Controls), the mean severity was 31.43 and the standard deviation 25.01. Respondents were considered to have a lifetime diagnosis if they met DSM-IV criteria for PTSD (measured using CAPS-IV) either current or past. Respondents were considered to have a current diagnosis if they met DSM-IV criteria for PTSD (measured using CAPS-IV) at the time of the assessment. The Institutional Review Board of the Minneapolis VA Health Care System approved this study.

### Detroit Neighborhood Health Study (DNHS, ADNH; Supplementary Table 1 #4 and #45)

See reference for details.^54^ Potentially traumatic events (PTEs) were identified using a list of 19 PTEs.^122^ The PTSD Checklist-Civilian (PCL-C) was used to assess PTSD over the lifetime by self-report during structured telephone interviews by referencing two traumatic events; one the respondent regarded as the worst and a second randomly selected event from the list of remaining PTEs (if the respondent experienced more than one traumatic event).^123^ Respondents were considered to have a diagnosis if all six DSM-IV criteria were met for either the worst or random event. Additional questions assessed the timing, duration, severity or illness, and disability resulting from symptoms. PTSD symptom severity, which ranged from 17 to 85, was assessed by summing the respondents’ ratings of the 17 post-traumatic symptoms on a scale indicating the degree to which the respondent was bothered by a particular symptom as a result of the worst trauma, ranging from 1 (not at all) to 5 (extremely). All DNHS participants, include N = 2081, of which cases N = 408 and controls N=1532. DNA for GWAS analysis was isolated from peripheral blood or saliva. The Institutional Review Board at the University of Michigan and University of North Carolina Chapel Hill approved this study.

### Drakenstein Child Health Study - South African sample (SAFR; Supplementary Table 1 #3)

See reference for details.^124 125 126^ The modified PTSD Symptom Scale (PSS) was used to assess PTSD.^127^ Specifically, the re-experiencing symptom cluster was considered met if the sum of reported symptoms totaled greater than or equal to 1; the avoidance/emotional numbing reported symptoms were greater than or equal to 3; and increased arousal cluster reported symptoms were greater than or equal to 2. Participants who scored above threshold in each of the clusters and had symptom duration for at least 1 month were classified as PTSD cases. The Faculty of Health Sciences human research ethics committee of the University of Cape Town (UCT) approved this study.

### EA CRASH (EACR; Supplementary Table 1 #42)

See reference for details.^128^ European American individuals were enrolled in the Emergency Department within 24 hours following motor vehicle collision (MVC) trauma/stress.^129^ The Impact of Events Scale-Revised (IES-R) was used to assess PTS symptom severity over the past week by research assistants 1 year following MVC.^129^ Respondents were considered to have a diagnosis if they scored 33 or higher on the IES-R questionnaire. The IES-R inventory calculates PTS symptom severity, which ranged from 0 to 88, by scoring a participant’s answers to 22 questions on a scale of 0-4 about symptoms of avoidance, intrusions, and hyperarousal. For this cohort (88 Cases, 276 Controls), the mean PTS symptom severity was 19.2 and the standard deviation was 18.4, measured 1-year after the MVC. DNA for GWAS analysis was isolated from blood collected in DNA PAXgene tubes. The Institutional Review Board of The University of North Carolina at Chapel Hill approved this study.

### Family Study of Cocaine Dependence and Collaborative Genetic Study of Nicotine Dependence (FSCD, COGA, COGB; Supplementary Table 1 #7-9)

See references for details.^130,131^ A module from the Diagnostic Interview Schedule for DSM-IV (DIS-IV),^132^ a structured assessment that evaluated the presence or absence of psychiatric disorders according to the DSM-IV^133^ criteria was used to evaluate PTSD in a sample of 471 cases and 3,568 controls. A history of fifteen specific traumatic events were queried including rape or sexual assault, assaultive violence (e.g., shot, stabbed), witnessing trauma to others, and non-violent trauma (e.g., serious accident, sudden death of a loved one). The traumatic events were assessed using closed-ended questions (e.g., Have you ever been raped or sexually assaulted?) with nominal response options (i.e., Yes or No). Participants were asked to select the most distressing event and were subsequently evaluated for symptoms of PTSD. A diagnosis of PTSD was dependent on Criterion A, which required intense fear, helplessness, or horror in association with the most distressing event. Interview data were checked for consistency by a senior editor and entered into a computerized data file. Lifetime psychiatric diagnoses were made by a computer algorithm that analyzed responses to the interview using DSM-IV criteria. The Washington University School of Medicine IRB approved the studies.

### Fort Campbell study (FTCB; Supplementary Table 1 #52)

Fort Campbell is a United States Army installation located astride the Kentucky-Tennessee border between Hopkinsville, Kentucky, and Clarksville, Tennessee. Fort Campbell is home to the 101st Airborne Division and the 160th Special Operations Aviation Regiment. One thousand, seven hundred and ninety-three (N=1793) active duty members of the Army’s 101st Airborne Division who deployed to Afghanistan participated in the study. Each participant was evaluated one-three times at the Fort Campbell U.S. Army military installation. The first evaluation took place prior to deployment in January-February, 2014, the second evaluation took place 3 days upon return from deployment and the third evaluation occurred 90-180 days upon return from deployment.

Potentially traumatic events were identified using self-report questionnaire that included PCL 5.^134^ The PTSD Checklist for DSM 5 (PCL-5) was used to assess PTSD during each phase of the study (pre-deployment, 3 days post deployment and 90-180 days post deployment).^134^ The PCL-5 score calculates PTSD symptom severity by summing the scores for all items, which ranged from 0 to 80 in the full range and ranged from 0 to 75 in this dataset. For this cohort (114 Cases, 1624 Controls), the mean severity was 6.81 and the standard deviation 11.24. Respondents were considered to have a current diagnosis if PCL5 score is at least 33. The Institutional Review Board of NYUMC and HARPO (DoD IRB) approved this study.

### Genetic and Environmental Predictors of Combat-Related PTSD (STRO; Supplementary Table 1 #35)

See reference for details.^135^ Subjects in this study were included from a larger STRONG STAR pre-/post-deployment study of deploying soldiers from Fort Hood in Killeen, Texas. The data included in this analysis are from the predeployment assessment. Potentially traumatic events were identified using the Life Events Checklist (LEC).^83^ The PTSD Checklist-Military Version (PCL-M) was used to assess PTSD over the prior month by self-repor.^76^ Subjects were considered to have a current diagnosis if the total score on the PCL was ≥ 50 and classified as a control if their PCL total score was < 50. In addition, both cases and controls were required to report having directly experienced or witnessing a traumatic event on the LEC. Based on these criteria, N=607 subjects were classified as having current PTSD and N=3,390 were classified as controls. DNA for GWAS analysis was isolated from blood. The Institutional Review Board of University of Texas Health Science Center at San Antonio approved this study.

### Genetic Risk for PTSD (YEHU; Supplementary Table 1 #56)

See reference for details.^136,137^ Potentially traumatic events were identified using the CAPS, SCID, MINI, and the Life Events Checklist.^81,83,138,139^ The CAPS, SCID, and MINI were used to assess PTSD during the past month or over the prior lifetime by a PhD level clinical psychologist.^81,138,139^ Respondents were considered to have a diagnosis if at least one Criterion B symptom, at least three Criterion C symptoms, at least two Criterion D symptoms, and Criterion A, E, and F (CAPS), J1, J2, and J3 are coded “yes”, at least three or more J4 questions are coded “yes”, at least two J5 answers are coded “yes”, and J6 is coded “yes” (MINI), and/or subject experienced a traumatic event and adverse consequences were experienced, both A criteria were coded 3, at least one B criteria was coded 3, at least three C criteria were coded 3, and at least two D criteria were coded 3 (SCID). Respondents were considered to have a current diagnosis if criteria for PTSD diagnosis were met within the past month. The CAPS calculates PTSD symptom severity, which ranged from 0 to 136. For this cohort (123 cases, 43 controls), the mean severity was 80.19 and the standard deviation 2.23. DNA for GWAS analysis was isolated from whole blood. The Institutional Review Board of James J. Peters VA Medical Center and Icahn School of Medicine at Mount Sinai approved this study.

### Genetics of Posttraumatic Stress Disorder/Substance Use Disorder Comorbidity (KSUD; Supplementary Table 1 #17)

See reference for details.^140^ Potentially traumatic events were identified using the Life Event Checklist for DSM-5 (LEC-5).^141^ The self-report PTSD Checklist for DSM-5 (PCL-5)^134^ was used to assess PTSD symptoms over the prior month. The PCL-5 contains 20 items which summed provide a measure of PTSD symptom severity (scores range from 0 to 80). For this cohort (137 Cases, 106 Controls), the mean severity was 40.57 and the standard deviation 20.96. Respondents were considered to have a current diagnosis if total symptom severity was at or above 38. The Institutional Review Board of Kent State University approved this study.

### GMRF-QUT (GMFR; Supplementary Table 1 #55)

See reference for details.^142^ Potentially traumatic events were identified using Criterion A event. The Clinician Administered PTSD Scale for DSM 5 (CAPS-5) was used to assess PTSD over the prior 2 weeks and lifetime by clinical psychologists.^143^

Respondents were considered to have a current diagnosis if CAPS-5 criteria were met. The CAPS-5 calculates PTSD symptom severity, which ranged from 0 to 56. For this cohort (100 Cases, 124 Controls), the mean severity was 9.63 and the standard deviation 10.05. DNA for GWAS analysis was isolated from peripheral blood. Ethics approval for the project was obtained from the Department of Veterans’ Affairs, Greenslopes Private Hospital, and Queensland University of Technology Human Research Ethics Committees. This study was carried out in accordance with The Code of Ethics of the World Medical Association (Declaration of Helsinki).

### Genetics Research and the Childbearing Year (GRAC; Supplementary Table 1 #54)

See reference for details.^144^ A total of 29 potentially traumatic events were identified using the Life Stressor Checklist^145^. PTSD symptoms were assessed using the National Women’s Study PTSD Module (NWS-PTSD), a widely used scale designed for use by lay interviewers, that consists of 20 items that assess DSM-IIl-R PTSD Criteria B, C, and D symptoms. The NWS-PTSD was performed by trained lay interviewers using computer-aided telephone interviewing and epidemiological methods (forced choice yes or no).^146,147^ Respondents were considered to have a diagnosis if lifetime DSM-IV PTSD criteria were met. Respondents were considered to have a current diagnosis if past month DSM-IV criteria were met. The NWS-PTSD assesses the number of PTSD symptoms endorsed, which ranged from 0 to 17. For this cohort (140 Cases, 138 Controls), the mean PTSD symptom count (out of 17 possible symptoms) was 6.4 and the standard deviation 5.5. DNA for GWAS analysis was isolated from saliva (Oragene tube). The Institutional Review Board of the University of Michigan approved this study.

### Grady Trauma Project (EGHS, GTPC; Supplementary Table 1 #44 and #47)

See reference for details.^148^ The modified PTSD Symptom Scale (PSS), a psychometrically valid 17-item self-report scale assessing PTSD symptomatology over the prior 2 weeks, was used to assess PTSD.^149^ Consistent with prior literature, the PSS frequency items (0 indicates not at all to 3 indicates ≥5 times a week) to obtain a continuous measure of PTSD symptom severity ranging from 0 to 51. For this sample, the PSS frequency items had standardized α=.90 (mean [SD], 13.81 [11.96]). No clearly established PSS cutoff score for PTSD diagnosis has been established; however, DSM-IV criteria for PTSD can be applied to PSS frequency items to create a proxy variable for PTSD diagnostic status. The Institutional Review Boards of Emory University School of Medicine and Grady Memorial Hospital approved this study.

### Growing Up Today Study (GUTS; Supplementary Table 1 #21)

See reference for details.^150^ Potentially traumatic events were identified using the Brief Trauma Questionnaire,^151^ plus questions on stalking and intimate partner violence (specific events queried included: witness an attack, get attacked, disaster, serious accident, attack on family member, stalked, family member killed in violence, served in war zone/saw war casualties, serious injury to self, physical intimate partner violence, sexual intimate partner violence). Breslau’s Short Screening Scale for DSM-IV PTSD was used to assess PTSD over the lifetime by self-report of symptoms.^152^ Respondents were considered to have a lifetime diagnosis if they reported experiencing 4 or more symptoms. Current diagnosis was not assessed. Breslau’s Short Screening Scale for DSM-IV PTSD calculates PTSD symptom severity, which ranged from 0 to 7, by counting the number of symptoms. For this cohort (312 Cases, 312 Controls), the severity was mean=2.63, SD=2.31 (cases: mean=4.93, SD=0.94; controls: mean=0.34, SD=0.47). DNA for GWAS analysis was isolated from saliva. The Institutional Review Board of Brigham and Women’s Hospital approved this study.

### Injury and Traumatic Stress Consortium (INTR; Supplementary Table 1 #27)

Subjects were participants in studies of the INTRuST Consortium, some of which are cited.^153–155^ Potentially traumatic events were identified using the Life Events Checklist for DSM-IV (LEC)^76^ and/or the Deployment Risk and Resilience inventory (DRRI).^156^ The PTSD Checklist for DSM-IV (PCL)^76^ was used to indicate a likely diagnosis of PTSD (or healthy control status); in some cases, this diagnosis was corroborated by the Clinician-Administered PTSD Scale (CAPS).^81,157^ The PCL was used as an indicator of PTSD symptom severity, which ranged from 17-85. DNA for GWAS analysis was isolated from whole blood. The Institutional Review Boards of UCSD (the Coordinating Center) and all the participating institutions approved this research.

### IVS (BRYA; Supplementary Table 1 #10)

See reference for details.^158^ Potentially traumatic events were identified using the Recent Life Events questionnaire.^79^ The Clinician Administered PTSD Scale (CAPS) was used to assess PTSD over the prior 4 weeks by psychologist interviewers.^81^ Respondents were considered to have a current diagnosis if DSM-IV were met. The CAPS calculates PTSD symptom severity, which ranged from 0 to 136, by clinical assessment. For this cohort (90 Cases, 312 Controls), the mean severity was 25.10 and the standard deviation 23.98. DNA for GWAS analysis was isolated from saliva. The Institutional Review Board of Western Sydney Area Health Service approved this study.

### Marine Resiliency Study (MRSC, BAKE; Supplementary Table 1 #1 and #57)

See reference for details.^10,159^ Participants were recruited from two studies including military personnel: (1) the Marine Resiliency Study, a prospective PTSD study with longitudinal follow-up (pre-and post-exposure to combat stress) of U.S. Marines bound for deployment to Iraq or Afghanistan, and (2) a crosssectional study involving a cohort of combat-exposed active duty or previously deployed service members (CAVC), including PTSD cases and controls with comparable psychosocial and clinical phenotypes. PTSD was diagnosed up to 3 times, once before deployment and 3 and/or 6 month post deployment. Posttraumatic stress (PTS) symptoms were assessed using a structured diagnostic interview, the Clinician Administered PTSD Scale (CAPS), and PTSD diagnosis followed the DSM-IV criteria.^81^ All participants included in this study met the *DSM-IV* criteria A1 event. For participants assessed at multiple timepoints, the timepoint with the highest CAPS score was used. Genomic DNA was prepared from blood leukocytes and genotyping was carried out by Illumina (http://www.illumina.com/) using the HumanOm-niExpressExome (HOEE) array with 951,117 loci and by RUCDR (http://www.rucdr.org) using the HOEE array with 967,537 loci. The study was approved by the University of California San Diego Institutional Review Board, and all participants pro-vided written informed consent to participate.

### McLean Trauma Sample (TEIC; Supplementary Table 1 #39)

The McLean Trauma Sample consists of three separate studies lead by Drs. Milissa Kaufman and Martin Teicher. See references for details.^160–162^ The first study was conducted at McLean Hospital’s Developmental Biopsychiatry Research Program (PI: Martin Teicher, MD, PhD) entitled “Sensitive Periods, Brain Development and Depression Study”. The group aimed to test the hypothesis that there are discrete sensitive periods when exposure to abuse or loss is maximally associated with risk for developing psychiatric disorders, specifically major depression and that risk for developing depression coincided with exposure to abuse during sensitive periods of hippocampal and prefrontal cortex vulnerability. These hypotheses were tested in a sample of 517 individuals (20-25 years of age) recruited from the community. Degree and timing of developmental exposure to abuse and loss across each childhood stage was quantified retrospectively using the Maltreatment and Abuse Chronology of Exposure (MACE) scale,^163^ as well as Traumatic Antecedent Interview,^164^ Childhood trauma Questionnaire^165^ and Adverse Childhood Experiences scale.^166^ Lifetime and current psychopathology including PTSD was assessed by trained psychologists, psychiatrists and clinical nurse specialists, using the Structured Clinical Interview for DSM-IV-TR.^167^ Respondents were considered to have a current diagnosis if DSM-IV-TR criteria were met in the preceding 30 days.

The second study was also conducted at McLean Hospital’s Developmental Biopsychiatry Research Program (PI: Martin Teicher). The key aims of the project were to test in a prospective study whether neurobiological correlates such as T2-relaxation time in dorsolateral prefrontal and anterior cingulate cortex and a large cerebellar lingual size can predict degree of drug and alcohol use in individuals with histories of childhood abuse and neglect, with an emphasis on sensitive periods of maximal exposure. These hypotheses were tested in a sample of 157 individuals (18-19 years of age) recruited from the community. Structured Clinical Interviews for DSM-IV (SCID-IV) Axis I and II psychiatric disorders were used for diagnoses.^167^ Mental health professionals (psychiatrists, Ph.D. psychologists, clinical nurse specialists) performed all the interviews and psychological assessments. In addition to the Maltreatment and Abuse Chronology of Exposure (MACE) scale,^163^ the 100-item semi-structured Traumatic Antecedents Interview^164^ was also used to assess maltreatment history, as well as the Childhood Trauma Questionnaire^165^ and the Adverse Childhood Experience score.^166^ PTSD was diagnosed using the SCID-IV-TR and the CAPS.^81^

The third study was conducted at McLean’s Dissociative Disorders and Trauma Research Program (PI: Milissa Kaufman, MD, PhD) entitled “Evaluating the Neurobiological Basis of Traumatic Dissociation in a Cross-Diagnostic Sample of Women with Histories of Childhood Abuse and Neglect”. Patients were recruited from inpatient and partial/residential treatment programs at McLean Hospital as part of a larger study on trauma-related dissociation comprised of diagnostic interviews, self-reports, neuropsychological testing, and neuroimaging protocols. All individuals endorsed a history of childhood trauma exposure, as assessed by the Traumatic Events Interview and Childhood Trauma Questionnaire. All participants also met criteria for DSM-5 PTSD as assessed by the Clinician-Administered PTSD Scale for DSM-5.

The Institutional review Board of McLean Hospital approved all studies. All saliva samples were collected using Oragene DNA collection kits (DNA Genotek) according to the instructions of the manufacturer.

### Mid-Atlantic Mental Illness Research Education and Clinical Center the study of Post-Deployment Mental Health Study (MIRE; Supplementary Table 1 #26)

See reference for details of the study of Post-Deployment Mental Health (1,308 cases, 1,914 controls).^168,169^ PTSD was diagnosed using the Structured Clinical Interview for DSM-IV Disorders (SCID) administered by trained interviewers.^82^ In accordance with the DSM-IV, PTSD consisted of three symptom clusters. These included re-experiencing symptoms (B symptoms), avoidance and numbing symptoms (C symptoms) and hyperarousal symptoms (D symptoms). Total PTSD symptoms and symptom clusters (B, C, or D) were measured using the Davidson Trauma Scale for all veterans, including individuals with current PTSD diagnosis and controls. The research was reviewed and approved by the Institutional Review Boards at the Salisbury, NC VA, Hampton, VA VA, Richmond, VAVA, Durham, NC VA and Duke University Medical Centers.

### National Centre for Mental Health (NCMH; Supplementary Table 1 #41)

See www.ncmh.info for details. Potentially traumatic events were identified by participant self-report. The Trauma Screening Questionnaire (TSQ) was used to assess PTSD over the prior 2 weeks by self-report.^170^ Respondents were considered to have a current diagnosis if a score of 6 or more was obtained. The TSQ calculates PTSD symptom severity, which ranged from 0 to 10, by selfreport. For this cohort (631 Cases, 653 Controls), the mean severity was 5.32 and the standard deviation 3.47. DNA for GWAS analysis was isolated from blood or saliva. The study was given a favorable ethical opinion by Wales Research Ethics Committee (REC) 2.

### National Health and Resilience in Veterans Study (NHRV; Supplementary Table 1 #13)

See reference for details.^171^ Potentially traumatic events were identified using the Trauma History Screen.^172^ The PTSD Checklist-Specific (PCL-S) was used to assess both lifetime and past-month PTSD symptoms related to respondent’s ‘worst’ traumatic event over their lifetimes by survey.^76^ Respondents were considered to have screen positive for PTSD if their PCL-S score was ≥50. The PCL-S calculates PTSD symptom severity, which ranged from 17 to 85, by selfreport. For this cohort (95 Cases, 1490 Controls), the mean PCL-S severity score was 26.9 and the standard deviation 11.3. DNA for GWAS analysis was isolated from saliva. The Human Subjects Subcommittee of VA Connecticut Healthcare System approved this study.

### NIU Trauma Study (NIUT; Supplementary Table 1 #40)

See reference for details.^173,174^ Potentially traumatic events were identified using the Traumatic Life Events Questionnaire and 12 questions regarding level of exposure to the campus shooting.^175^ The Distressing Events Questionnaire was used to assess PTSD immediately following the mass shooting (average 3.2 weeks) by self-report.^176^ Respondents were considered to have a diagnosis if their DEQ score was ≥ 18. The DEQ calculates PTSD symptom severity, which ranged from 0 to 66, by self-report. For this cohort (280 Cases, 411 Controls), the mean severity was 16.49 and the standard deviation 12.35. DNA for GWAS analysis was obtained from 204 of the PTSD cases and was isolated from saliva. The Institutional Review Board of Northern Illinois University approved this study.

### Nurses Health Study II (NHS2; Supplementary Table 1 #5)

See reference for details.^177^ Participants identified stressful events they had experienced from a list of 25 events used in diagnostic interviews, ^178, 102,179–181^ and PTSD was assessed in relation to participants’ self-selected worst stressful event. Participants were cued to think of the period following the event during which symptoms were most frequent and intense. They were asked whether they had ever been bothered by each of 17 symptoms and rated each symptom on a Likert-style scale (1: “not at all” to “5: extremely”).^182^ Additional questions assessed the other three DSM-IV criteria: intense fear, horror, or helplessness in response to the event (Criterion A2), symptom duration of at least one month (Criterion E), and clinically significant impairment in functioning due to symptoms (Criterion F)^178^. Based on the diagnostic interview, we created two lifetime PTSD phenotypes as follows.

To meet criteria for lifetime PTSD diagnosis, respondents must have endorsed experiencing one or more of the 5 re-experiencing symptoms, 3 or more of the 7 avoidance/numbing symptoms, 2 or more of the 5 arousal symptoms, and criteria A2, E and F as defined above. In addition to the diagnostic phenotype, we analyzed lifetime PTSD symptom severity which was defined as the sum of the symptom ratings across the 17 questions.

The reliability of the PTSD diagnosis was assessed using a blind review of audiotapes from 50 interviews and the Cohen’s kappa statistic was 1.0 (perfect reliability).^183^ We assessed the validity of our identification of PTSD in a separate cohort, the Detroit Neighborhood Health Study, via clinical interviews among a random subsample of 51 participants and found excellent concordance.^54^ The Partners Human Research Committee approved this study.

### Nurses Health Study II (NHSY; Supplementary Table 1 #22)

See reference for details.^184^ The Nurses’ Health Study II is an ongoing cohort of 116,430 female nurses initially enrolled in 1989 and followed with biennial questionnaires. The present study included follow-up through 2013. This study included women who returned a supplementary 2008 questionnaire on trauma exposure and PTSD symptoms (N=54,763). This questionnaire was sent to a subsample of participants (N=60,804, response rate=90.1%). To retain participation in the ongoing longitudinal cohort, only women who have already returned their biennial questionnaire are sent supplementary questionnaires. Women missing data on trauma or PTSD symptoms (N=3,930) were excluded. This study was approved by the Institutional Review Board of Brigham and Women’s Hospital. Return of the questionnaire via US mail constitutes implied consent.

Trauma exposure and PTSD symptoms were assessed on a supplementary 2008 questionnaire. The 16-item Brief Trauma Questionnaire queried lifetime exposure to 15 types of traumatic events (e.g., serious car accident, sexual assault) and an additional item queried any traumatic event not covered in the other questions. ^151^ Respondents were asked to identify which trauma was their worst or most distressing; they were then asked their age at this worst trauma as well as their age at their first trauma. PTSD symptoms were assessed in relation to their worst trauma with the 7-item Short Screening Scale for DSM-IV PTSD.^185^ Four or more symptoms on this scale have been associated with PTSD diagnosis (sensitivity=80%, specificity=97%, positive predictive value=71%, negative predictive value=98%).^185^ For cases women with ≥4 PTSD symptoms were selected, for controls women with <4 PTSD symptoms (most had 0) were selected.

### Ohio National Guard (ONGA; Supplementary Table 1 #2)

See reference for details.^42^ A total of 37 potentially traumatic events were identified using the Clinician-Administered PTSD Scale (CAPS-IV)^81^ and the 1996 Detroit Area Survey of Trauma.^122^ PTSD symptoms were assessed using a 17-item structured interview scale derived from the PTSD Checklist (PCL) for DSM-IV^76^ performed by trained lay telephone interviewers using epidemiological methods (forced choice symptom severity range, 1-5). Reliability of the telephone interview was validated against the criterion standard (in-person CAPS interview by mental health professional) in a clinical subsample (N = 500), demonstrating high specificity (0.92).^186^ Respondents were considered to have a diagnosis if lifetime DSM-IV PTSD criteria were met. Respondents were considered to have a current diagnosis if past month DSM-IV criteria were met. The PCL calculates PTSD symptom severity, which ranged from 17 to 85, by sum of scores of items endorsed. For this cohort (125 Cases, 125 Controls), the mean severity was 38.4 and the standard deviation 17.6. DNA for GWAS analysis was isolated from saliva (Oragene tube). The Institutional Review Board of VA Ann Arbor Health System approved this study.

### OPT (FEEN; Supplementary Table 1 #37)

Potentially traumatic events were identified using the standard trauma interview.^187^ The PSS-I was used to assess PTSD over the prior two weeks for the trauma of interest by postdoctoral and graduate level assessors trained to reliability.^105^ The SCID-IV was used to assess lifetime PTSD (not current) for a single trauma not the focus of treatment.^82^ Respondents were considered to have a current PTSD diagnosis if they met DSM-IV diagnostic criteria and had a PSS-I score of 25 or greater. The PSS-I also provides PTSD symptom severity, with a range from 0 to 51. For this cohort (118 cases), the mean severity was 30.36 and the standard deviation 6.91. DNA for GWAS analysis was isolated from blood. The Institutional Review Board of University Hospitals approved this study.

### Portugal (PORT; Supplementary Table 1 #20)

See reference for details.^188^ Potentially traumatic events were identified using CIDI, which included a module on DSM-IV PTSD that inquired about lifetime exposure to each of 27 different traumatic events (criterion A1). Respondents who reported ever experiencing any of the traumatic events were then asked about the number of exposures (NOE) and age at first exposure (AOE) for each.^178^ The CIDI was used to assess PTSD over the prior Lifetime DSM-IV PTSD and other common DSM-IV disorders.^178^ DNA isolated from saliva. The Institutional Review Board of the Nova Medical School, Portugal, approved this study.

### PRISMO (PRIS; Supplementary Table 1 #25)

See reference for details.^189^ Potentially traumatic events were identified using a 19-item checklist.^189^ The Dutch Self-Rating Inventory for PTSD (SRIP) was used to assess PTSD one month prior to deployment and up to 2 years after deployment.^190^ Respondents were considered to have a current diagnosis if a SRIP total score of ≥ 38 at any measurement was met. The SRIP calculates PTSD symptom severity, which ranged from 22 to 88, by self-report. For this cohort (144 Cases, 815 Controls), the mean severity was 27.30 and the standard deviation 6.38. DNA for GWAS analysis was isolated from whole blood. The Institutional Review Board of Utrecht University Medical Center approved this study.

### Pregnancy Outcomes, Maternal and Infant Study Cohort (PROM; Supplementary Table 1 #23)

See reference for details.^191^ Potentially traumatic events were experiences of intimate partner violence [identified using Demographic Health Survey Questionnaires and Modules: Domestic Violence Module^192^ and the World Health Organization Multi-Country Study on Violence Against Women]^193^ and history of childhood abuse [identified using the Childhood Physical and Sexual Abuse Questionnaire adopted from the Center for Disease Control and Prevention Adverse Childhood Experiences Study].^194^ The Posttraumatic Stress Disorder Checklist – Civilian Version (PCL-C) was used to assess PTSD over the prior month.^195^ The PCL-C is a self-report measure with 17 items reflecting DSM-IV symptoms of PTSD and closely follows the Diagnostic and Statistical Manual of Mental Disorders (DSM-IV) criteria. For each item, participants were asked how bothered they were by a symptom over the past month on a 5-point Likert scale ranging from 1: “not at all” to 5: “extremely” in regards to their most significant life event stressor. The total score on the PCL-C ranges from 17 to 85. Recent data from our team support that a PCL-C score of 26 or higher on the Spanish-language version, is associated with an 86% sensitivity and 63% specificity in diagnosing PTSD in a Peruvian population using the Diagnostic and Statistical Manual of Mental Disorders (DSM-IV) criteria.^195^

All study procedures were approved by the Instituto Nacional Materno Perinatal in Lima, Peru, and the Office of Human Research Administration at the Harvard T.H. Chan School of Public Health, Boston, MA.

### Readiness and Resilience in National Guard Soldiers (RING; Supplementary Table 1 #33)

See reference for details.^196^ Potentially traumatic events were identified using the Clinician-Administered PTSD Scale for DSM-IV (CAPS-IV) and items from the Deployment Risk and Resilience Inventory (DRRI)^81,121^ The CAPS-IV and the PTSD Checklist (PCL) were also used to assess PTSD currently (over the prior 2 weeks) and over the lifetime by trained interviewers and through self-report.^81,197^ The PCL calculates PTSD symptom severity, which ranged from 17 to 73 currently and 17 to 85 lifetime, by self-report. For this cohort (41 Cases, 162 Controls), the mean current severity was 34.03 and the standard deviation was 13.48. The mean lifetime severity was 33.89 and the standard deviation was 12.98. Respondents were considered to have a lifetime diagnosis if they had a PCL score ≥ 50 at the one-year post-deployment time point. Respondents were considered to have a current diagnosis if the same criteria were met at the two-year post-deployment time point. The Institutional Review Board of the Minneapolis VA Health Care System approved this study.

### Risbrough/Norman randomized controlled psychotherapy trial (VRIS; Supplementary Table 1 #58)

See reference for details.^198^ Potentially traumatic events were identified using the life event checklist. The CAPS was used to assess PTSD over the prior over the past month by interviewers.^143^ Respondents were considered to have a diagnosis if DSM-5 diagnostic criteria using the CAPS-5 (a criterion A trauma and 2 or higher on 1 re-experiencing symptom (criterion b), 1 avoidance symptom (criterion c), 2 negative alterations in cognitions or mood (criterion d), and 2 hyperarousal symptoms (criterion e)) were met. The CAPS calculates PTSD symptom severity, which ranged from 0 to 80 by clinician rater of symptoms and severity. For this cohort (73 Cases, 5 sub-clinical), the mean severity was 42 and the standard deviation 9.6. DNA for GWAS analysis was isolated from saliva using Oragene. The Institutional Review Board of The San Diego VA approved this study.

### Shared Roots (SHRS; Supplementary Table 1 #46)

See reference for details.^141^ Potentially traumatic events were identified using The Life Events Checklist for DSM-5 (LEC-5).^141^ The Clinician-Administered PTSD Scale for DSM-5 (CAPS-5)^143^ was administered by clinicians to assess PTSD over the prior month. The CAPS-5 and the PTSD Checklist for DSM-5 (PCL-5)^134^ calculates PTSD symptom severity, with a score range of 0 to 80, by adding scores ranging from 0 to 4 for all twenty items. For this cohort (164 Cases, 164 Controls), the mean severity on the PCL-5 was 33.0 and the standard deviation 23.9. A lifetime diagnosis of PTSD was not assessed for in this study. Respondents were considered to have a current diagnosis if DSM-5^199^ criteria based on the CAPS-5 were met. The Institutional Review Board of Stellenbosch University approved this study.

### Southeastern Europe PTSD (SEEP; Supplementary Table 1 #49)

See reference for details.^200^ Potentially traumatic events were identified using Life Stressor List and List of traumatic events including frequency and severity of traumatic events.^201^ The CAPS was used to assess lifetime PTSD by medical personnel (psychiatrists, psychologists or psychiatric residents).^81^ Respondents were considered to have a diagnosis if DSM-IV criteria were met over lifetime. The CAPS calculates PTSD symptom severity, which ranged from 27 to 141. For this cohort (347 Cases, 339 Controls), the mean severity was 74.3 and the standard deviation 20.2. DNA for GWAS analysis was isolated from EDTA blood. The Institutional Review Board of the universities of Sarajevo, Zagreb, Tuzla, Mostar, Prishtina and Würzburg approved this study.

### Study of Aftereffects of Trauma: Understanding Response in National Guard (SATU; Supplementary Table 1 #11)

See reference for details.^202^ Potentially traumatic events were identified using Clinician-Administered PTSD Scale for DSM-IV (CAPS-IV).^81^ The CAPS-IV was also used to assess PTSD currently (over the prior 2 weeks) and over the lifetime by trained interviewers^81^. The CAPS-IV calculates PTSD symptom severity, which ranged from 0 to 105. For this cohort (88 Cases, 62 Controls), the mean severity was 50.33 and the standard deviation 24.55. Respondents were considered to have a lifetime diagnosis if they met DSM-IV criteria for PTSD (measured using CAPS-IV) either current or past. Respondents were considered to have a current diagnosis if they met DSM-IV criteria for PTSD (measured using CAPS-IV) at the time of the assessment. The Institutional Review Board of the Minneapolis VA Health Care System approved this study.

### Sydney Neuroimaging (BRY2; Supplementary Table 1 #36)

Potentially traumatic events were identified using clinical interview to identify history of traumatic events. The CAPS was used to assess PTSD over the prior 4 weeks by Masters level clinical interviewers.^81^ Respondents were considered to have a diagnosis if DSM-IV criteria were met. The CAPS calculates PTSD symptom severity, which ranged from 0 to 136, by clinical interview. For this cohort (82 Cases, 86 Controls), the mean severity was 39.67 and the standard deviation 31.75. DNA for GWAS analysis was isolated from saliva. The Institutional Review Board of Western Sydney Area Health Service approved this study.

### VA Boston-National Center for PTSD Study (NCPT, TRAC; Supplementary Table 1 #31, Supplementary Table 1 #32)

A total of 437 white non-Hispanic cases and 215 trauma-exposed controls is the composite of two datasets. The first from a cohort of white non-Hispanic subjects as described in a previous GWAS^8^ that passed ancestry filters as performed using SNPweights^203^ according to PGC-PTSD protocols (300 cases and 165 controls; 305 males and 160 females). The majority of this sample consisted of US veterans, but also included the intimate partners of a subset of the veterans. The second cohort was made up of subjects from the Translational Research Center for TBI and Stress Disorders, a VA RR&D Traumatic Brain Injury Center of Excellence at VA Boston Healthcare System (TRACTS) study of US veterans. From TRACTS, 137 white non-Hispanic cases and 50 controls passed ancestry filters based on SNPweights and were included in the analysis. The TRACTS sample is largely male (170 men and 17 women). The genotyping, quality control, filtering and imputation for these cohorts has been described in detail elsewhere.^8,204^ Briefly, genotyping was performed using the Illumina HumanOmni2.5-8 microarrays (Illumina, San Diego, CA). Imputation of non-genotyped SNPs was performed using IMPUTE2^205–208^ and 1000 genomes phase 1 reference data.^209^ Principal components were generated by the program EIGENSTRAT^210^ based on 100,000 SNPs. For both cohorts, participants were administered the Clinician-administered PTSD scale for DSM-IV (CAPS-IV),^81^ a 30-item structured diagnostic interview that assesses the frequency and severity of the 17 DSM-IV PTSD symptoms, 5 associated features and functional impairment, and both current and lifetime PTSD symptoms. These studies were performed under the oversight of the appropriate VA health care facilities institutional review boards.

### UK Biobank Cohort Description for PGC-PTSD (UKBB; Supplementary Table 1 #60)

The UK Biobank is an epidemiological resource assessing a range of health-related phenotypes in approximately 500,000 British individuals who were recruited between the ages of 40 and 70.^211^ Genome-wide genotype data is available on all participants, as well as a broad range of health phenotypes assessed at varying intensity. Data from an online follow-up questionnaire assessing common mental health traits, including questions designed to screen for PTSD, was available on 157,366 individuals.^16^

#### Phenotype

PTSD phenotypes were derived from the mental health online follow-up of the UK Biobank (Resource 22 on http://biobank.ctsu.ox.ac.uk). Participants were asked six questions derived from the brief civilian version of the PTSD Checklist Screener (PCL-S;^157^) assessing PTSD symptoms in the prior month. Questions comprised three initial questions related to avoidance of activities, disturbing thoughts, and feeling upset, and three further questions related to feeling distant, feeling irritable and having trouble concentrating (UK Biobank fields 20494-20498, 20508). Each item was scored on a five-point scale according to the amount of concern caused by that item in the past month (1=“Not at all” to 5=“Extremely”). The final item concerning trouble concentrating was drawn from an equivalent item from the Patient Health Questionnaire-9 (PHQ9) depression questionnaire and was scored on a four-point scale according to frequency of difficulties (1=“Not at all” and 4=“Nearly every day”). All items were summed for each individual to yield a total score ranging 3-29. Cases were defined as all individuals with a PCL-S score ≥ 13, controls as all individuals who responded to all of the initial three questions and had PCL-S score ≤ 12.

Two sensitivity analyses were performed to assess the stability of the UK Biobank PTSD phenotype. Firstly, controls were limited only to those who answered all initial questions with “Not at all” and so had a PCL-S score = 3. Secondly, PTSD cases and controls were limited to those reporting a lifetime trauma exposure. To assess the impact of reported trauma exposure on the PTSD phenotype, a trauma exposure measure was derived from questions in the mental health online follow-up that related to common triggers of post-traumatic stress-disorder.^16^ These questions asked if participants had ever: experienced combat; had a life-threatening accident; been diagnosed with a life-threatening illness; been a victim of a physically violent crime; been a victim of sexual assault; or witnessed a sudden violent death. Responses were combined to a single variable capturing any report of traumatic experience versus no report.

#### Genetic quality control

Genetic data for analyses was obtained from the full release of the UK Biobank data (N=487,410).^212^ Individuals were removed if this was recommended by the UK Biobank for unusual levels of missingness or heterozygosity; if call rate < 98% on genotyped SNPs; if they were related to another individual in the dataset (KING r < 0.044, equivalent to removing up to third-degree relatives inclusive); and if the phenotypic and genotypic gender information was discordant (X-chromosome homozygosity (FX) < 0.9 for phenotypic males, FX > 0.5 for phenotypic females). Removal of relatives was performed using a greedy algorithm, which minimise exclusions (for example, by excluding the child in a mother-father-child trio). All analyses were limited to individuals of White Western European ancestry, as defined by 4-means clustering on the first two genetic principal components provided by the UK Biobank.^213^ Principal components analysis was also performed on the European-only subset of the data using the software package flashpca2^214^. After quality control, individuals were excluded from analysis if they did not complete the mental health online questionnaire (N=126,522).

Genetic analyses used imputed variants provided by the UK Biobank.^212^ Autosomal genotype data from two highly-overlapping custom genotyping arrays (covering ∼800,000 markers) underwent centralised quality control to remove genotyping errors before being imputed in a two-stage imputation to the Haplotype Reference Consortium (HRC) and UK10K (for rarer variants not present in the HRC) reference panels ^(215;216;212)^. In addition, this central quality control, variants for analysis were limited to common variants (minor allele frequency > 0.01) imputed with higher confidence (IMPUTE INFO metric > 0.4). In addition, only variants that were directly genotyped or that were imputed from the HRC were included.^215^

Genome-wide association analyses: Prior to analysis, PTSD status was residualised on six prinicipal components from the genetic data and factors capturing genotyping batch and recruitment centre, using logistic regression. GWAS were then performed on the resulting deviance residuals using linear regressions on imputed genotype dosages in BGenie v1.2, software written for genetic analyses of UK Biobank.^212^

### Vietnam Era Twin Study of Aging (VETS; Supplementary Table 1 #24)

See references for details.^217,218^ Potentially traumatic events were identified using the Combat Exposure Index^219^ and the Diagnostic Interview Schedule Version III-Revised (DIS-III-R).^dis-iii-R; 220^ The Vietnam Era Twin Registry PTSD scale^221^ (administered at average age 38) was used to assess PTSD symptoms over the past 6 months and the PCL-civilian version for DSM-IV^222^ (administered at average age 62) was used to assess PTSD over the prior month. These two 223 instruments correlate 0.90 when administered at the same time.^223^ The 17-item PCL calculated PTSD symptom severity, which ranged from 17 to 84. Each response was rated on a 1-5 scale (from “not at all” to “extremely”). For this cohort (60 Cases, 841 Controls), the mean severity was 26.2 and the standard deviation was10.5. For this analysis, respondents were considered to have a diagnosis of PTSD if they met DSM-III-R criteria based on the DIS-II-R interview. DNA for GWAS analysis was isolated from blood. Genotyping was performed by deCODE Genetics, Reykjavik, Iceland. The Institutional Review Boards of the University of California, Sand Diego, Boston University, and the Puget Sound VA Healthcare System approved this study.

### The Women and Children’s Health Study (WACH; Supplementary Table 1 #43)

See reference for details.^224^ Potentially traumatic events were identified using the Life Events Checklist (LEC) for DSM-5.^141^ The PTSD Checklist for DSM-5 (PCL-5) was used to assess PTSD over the lifetime by interviewers.^225^ Respondents were considered to have a diagnosis of PTSD if the PCL-5 score was >=38. Respondents were considered to have a current diagnosis based only upon self-report during the interview. The PCL-5 calculates PTSD symptom severity, which ranged from 0 to 79. For this cohort (151 Cases, 150 Controls), the mean severity was 52.4 (SD: 11.2) for cases and 31.0 (SD: 23.5) for controls DNA for GWAS analysis was isolated from blood. The Institutional Review Board of the Louisiana State University Health Sciences Center-New Orleans approved this study.

### Yale-Penn Study (GSDC; Supplementary Table 1 #6)

See reference for details.^226^ Sample collection and diagnostic interviews were performed by trained interviewers using the Semi-Structured Assessment for Drug Dependence and Alcoholism (SSADDA; available at https://zork.wustl.edu/nida/studydescriptions/study1/ssaddav112ns.pdf) to derive diagnoses for lifetime psychiatric and substance use disorders based on DSM-IV criteria. Twelve types of traumatic events were assessed: experienced direct combat in a war; seriously physically attacked or assaulted; physically abused as a child; seriously neglected as a child; raped; sexually molested or assaulted; threatened with a weapon; held captive or kidnapped; witnessed someone being badly injured or killed; involved in a flood, fire, or other natural disaster; involved in a life-threatening accident; suffered a great shock because one of the above events happened to someone close to you; and other. Participants were asked to list up to three traumatic events and describe the trauma in detail. Those reporting traumatic experiences were then interviewed for potential PTSD symptoms. After the data were scored, PTSD diagnoses were generated based on DSM-IV criteria. The institutional review boards at Yale University School of Medicine, the University of Connecticut Health Center, the University of Pennsylvania School of Medicine, the Medical University of South Carolina, and McLean Hospital approved the study.

### Data assimilation

Subjects were genotyped on a range of Illumina genotyping arrays (exception: UKBB was genotyped on the Affymetrix Axiom array). At the time of analysis, direct access to individual-level genotypes was permitted for 65,555 subjects. For these, pre-QC’ed genotype data was deposited on the LISA server for central data processing and analysis, using the standard PGC pipelines (https://sites.google.com/a/broadinstitute.org/ricopili/ and https://github.com/orgs/Nealelab/teams/ricopili). Studies with data sharing restrictions (8 studies, N = 137,114 subjects) performed analyses off site using identical pipelines unless otherwise indicated (**Supplementary Table 1**). Such studies then shared summary results for meta-analyses.

### Global Ancestry Determination

To determine consistent global ancestry estimates across studies, each subject was run through a standardized pipeline, based on SNPweights^203^ of 10,000 ancestry informative markers genotyped in a reference panel including 2,911 unique subjects from 71 diverse populations and 6 continental groups (K=6)^227^ (https://github.com/nievergeltlab/global_ancestry). Pre-QC genotypes were used for these analyses.

For the present GWA studies, subjects were placed into 3 large, homogeneous groupings, using previously established cut-offs (**Supplementary Table 2**): European and European Americans (EUA; subjects with ≥90% European ancestry), African and African-Americans (AFA; subjects with ≥5% African ancestry, <90% European ancestry, <5% East Asian, Native American, Oceanian, and Central-South Asian ancestry; and subjects with ≥ 50% African ancestry, <5% Native American, Oceanian, and <1% Asian ancestry), and Latinos (AMA; subjects with ≥5% Native American ancestry, <90% European, <5% African, East Asian, Oceanian, and Central-South Asian ancestry). Native Americans (subjects with ≥60% Native American ancestry, <20% East Asian, <15% Central-South Asian, and <5% African and Oceanian ancestry) were grouped together with AMA. All other subjects were excluded from the current analyses (N=6,740). **Supplementary Figure 1** shows the ancestry grouping used for GWAS of 69,484 subjects for which individual-level genotype data was available to the PGC. The ancestry pipeline was shared with external sites in order to ensure consistency in ancestry calling across cohorts.

### Genotype quality control

The standard PGC pipeline RICOPILI was used to perform QC, but modifications were made to allow for ancestrally diverse data. In the modified pipeline, each dataset was processed separately, including subjects of all ancestries. Sample exclusion criteria: using SNPs with call rates >95%, samples were excluded with call rates <98%, deviation from expected inbreeding coefficient (fhet < −0.2 or > 0.2), or a sex discrepancy between reported and estimated sex based on inbreeding coefficients calculated from SNPs on X chromosomes. Marker exclusion criteria: SNPs were excluded for call rates <98%, a >2% difference in missing genotypes between cases and controls, or being monomorphic. Hardy-Weinberg equilibrium (HWE): the modified pipeline identified the largest ancestry group in the data, identified SNPs with a HWE p-value <1×10^−6^ in controls, and excluded these SNPs in all subjects of the specific datasets, irrespective of ancestry.

### Relatedness within studies

Within-study relatedness was estimated using the IBS function in PLINK 1.9.^228^ From each pair with relatedness 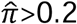, one individual was removed from further analysis, retaining cases where possible.

### Calculation of principal components (PC’s) for GWAS

For each dataset, unrelated subjects were subset into the 3 ancestry groups (EUA, AFA, AMA; **Supplementary Tables 3-5**) for analysis. SNPs were excluded that had a MAF <5%, HWE p >1×10^−3^, call rate <98%, were ambiguous (A/T, G/C), or due to being located in the MHC region (chr. 6, 25-35 MB) or chromosome 8 inversion (chr. 8, 7-13 MB). SNPs were pairwise LD-pruned (r^2^ >0.2) and a random 100K markers were used to calculate 3 sets of PC’s based on the smartPCA algorithm in EIGENSTRAT.^210^

### Imputation

Imputation was based on the 1000 Genomes phase 3 data (1KGP phase 3^229^). Any dataset using a human genome assembly version prior to GRCh37 (hg19) was lifted over to GRCh37 (hg19). SNP alignment proceeded as follows: for each dataset, SNPs were aligned to the same strand as the 1KGP phase 3 data. For ambiguous markers, the largest ancestry group was used to calculate allele frequencies and only SNPs with MAF<40% and ≤15% difference between matching 1KGP phase 3 ancestry data were retained. Pre-phasing was performed using default settings in SHAPEIT2 v2.r83 7 ^230^ without reference subjects, and phasing was done in 3 megabase (MB) blocks, where an additional

1 MB of buffer was added to either end of the block. Haplotypes were then imputed using default settings in IMPUTE2 v2.2.2,^205^ with 1KGP phase 3 reference data and genetic map, a 1 MB buffer, and effective population size set to 20,000. RICOPILI default filters for MAF and Info were removed since analyses were run across ancestry groups at this step. Imputed datasets were deposited with the PGC DAC and are available for approved requests.

### Main GWAS

The analysis strategy for the main association analyses is shown in **Supplementary Tables 3-5**. Analyses were performed separately for each study and ancestry group, unless otherwise indicated. The minimum number of subjects per analysis unit was set at 50 cases and 50 controls, or a total of at least 200 subjects, and subsets of smaller size were excluded. Smaller studies of similar composition were genotyped jointly in preparation for joint analyses. For studies with unrelated subjects, imputed SNP dosages were tested for association with PTSD under an additive model using logistic regression in PLINK 1.9, including the first five PC’s as covariates. For family and twin studies (VETSA, QIMR), analyses were performed using linear mixed models in GEMMA v0.96 ^231^, including a genetic relatedness matrix (GRM) as a random effect to account for population structure and relatedness, and the first five PC’s as covariates. The UKBB data (UKBB) were analyzed with BGenie v1.2 (https://www.biorxiv.org/content/early/2017/07/20/166298) using a linear regression with 6 PCs, and batch and center indicator variables as covariates (see cohort description for details). In addition, all GWAS analyses were also performed stratified by sex.

### Meta analyses

Summary statistics on the linear scale (from GEMMA and BGenie) were converted to a logistic scale prior to meta-analysis (for formula see ^232^). Within each dataset and ancestry group, summary statistics were filtered to MAF ≥1% and PLINK INFO score ≥0.6. Meta-analyses across studies were performed within each of the 3 ancestry groups and across all ancestry groups. Inverse variance weighted fixed effects meta-analysis was performed with METAL (v. March2 5 2 011).^233^ Heterogeneity between datasets was tested with a Cochran test and for nominally significant Q-values, a Han-Eskin random effects model (RE-HE) meta-analysis was performed with METASOFT v.2.0.1.^234^ Markers with summary statistics in less than 25% of the total effective sample size or present in less than 3 studies were removed from meta-analyses. Quantile-quantile (QQ) plot of expected versus observed −log_10_ p-values included imputed and genotyped SNPs at MAF ≥ 1%.

For genome-wide significant hits, Forest plots and PM-Plots (-log_10_(P-values vs. M-values)) were generated using the programs METASOFT with default settings and M-values were generated using the MCMC option.^17,235^ Regional association plots were generated using LocusZoom^236^ with 400KB windows around the index variant and compared to the corresponding windows in the other ancestry groups, including the 1000 Genomes Nov. 2014 reference populations EUR, AFR, and AMR, respectively. To test for sex-specific effects, a z-test was performed on the difference of the effect estimates from male and female sex-stratified analyses.

### Comparability of PGC2 studies

To evaluate the comparability of the 60 different studies, each EUA GWAS was assigned at random to one of two groups using a biased coin design (**Supplementary Figure 2**). A Bayesian rule was used to set the level of bias^237^, where the sample size parameters were set to the effective sample size (4/(1/N cases + 1/N controls)) of each group and gamma was set to 5×10^−5^. After all datasets were assigned to a group, summary statistics of GWAS were inverse variance weighted meta-analyzed in METAL^233^. Genetic correlations (*r_g_*) between groups were calculated in LDSC^15^ using meta-analysis summary statistics. Ten repetitions of random assignment and subsequent *r_g_* calculation were performed.

Similarly, LDSC was used to calculate genetic correlation between UKBB and a GWAS meta-analysis including all Freeze 1.5 EUA subjects, between male and female subsets, as well as between different phenotype selections in UKBB.

### Local ancestry deconvolution

A pipeline was developed to determine local ancestry in subjects with African and/or Native American admixture (AFA, AMA; **Supplementary Figure 3**). Additional QC to consistently prepare cohort data for downstream analysis was performed with a custom script (https://github.com/eatkinson/Post-QC). Post-QC steps involved extracting autosomal data, removing duplicate loci, updating SNP IDs to dbSNP 144,^238^ orienting data to the 1KGP reference (with removal of indels and loci that either were not found in 1KGP or that had different coding alleles), flipping alleles that were on the wrong strand, and removing ambiguous SNPs.

#### Data harmonization and phasing

We then intersected and jointly phased the post-QC’ed cohort data with autosomal data from 247 1KGP reference panel individuals, removing conflicting sites and flipping any remaining strand flips. The merged dataset was then filtered to include only informative SNPs present in both the cohort and reference panel using a filter of MAF≥0.05 and a genotype missingness cutoff of 90%. The program SHAPEIT2^239^ was used to phase chromosomes, informed by the HapMap combined b37 recombination map.^240^ Individuals from the cohort and reference panel were then separated and exported as harmonized sample and reference panel VCFs to be fed into Rfmix.^241^

#### Reference panel

Three ancestral populations of European, African, and Native American ancestry were chosen for the admixed AFA cohorts based on ancestry proportion estimates from SNPweights runs. All reference populations were taken from 1KGP phase 3 data.^242^ Specifically, 108 West African Bantu-speaking YRI were used as the African reference population, 99 CEU comprised the European reference, and 40 PEL of >85% Native American ancestry were used as the Native American reference panel. Individuals used as the reference panel can be found on (https://github.com/eatkinson).

#### Local ancestry inference (LAI) parameters

LAI was run on each cohort separately using RFMix version 2^241^ (https://github.com/slowkoni/rfmix) with 1 EM iteration and a window size of 0.2 cM. We used the HapMap b37 recombination map^240^ to inform switches. The −n 5 flag (terminal node size for random forest trees) was included to account for an unequal number of reference individuals per reference population. We additionally used the --reanalyze-reference flag, which recalculates admixture in the reference samples for improved ability to distinguish ancestries.

#### Local ancestry of genome-wide significant variants

Haplotypes of the genomic regions around genome-wide significant associations were aligned to the local ancestry calls according to genomic position. To compare MAF of top hits on different ancestral backgrounds within a specific admixed population (AFA or AMA), subjects were grouped according to the number of copies (1 or 2) of a specific ancestry (European, African, and Native American, respectively) at that position. For a given SNP, MAF was calculated within each of the 6 groups. To ensure successful elimination of population stratification by standard global PC’s in regression analyses of admixed populations, two (out of 3, to reduce redundancy) local ancestry dosage covariates were included, coded as the number of copies (0, 1, or 2) from a given ancestral background. Finally, to compare if effects of the minor allele depend on a specific ancestral background (European, African, and Native American), for each SNP, we coded variables that counted the number of copies of the minor allele per ancestral background. Association between these three variables and PTSD were jointly evaluated using a logistic regression, including study indicators and five global ancestry PC’s as additional covariates.

### Functional mapping and annotation

We used FUnctional Mapping and Annotation of genetic associations (FUMA) v1.3.0 (http://fuma.ctglab.nl/) to annotate GWAS data and obtain functional characterization of risk loci ^243^. Annotations are based on human genome assembly GRCh37 (hg19). FUMA was used with default settings unless stated otherwise. The SNP2Gene module was used to define independent genomic risk loci and variants in LD with lead SNPs (r^2^ > 0.6, calculated using ancestry appropriate 1KGP reference genotypes). SNPs in risk loci were mapped to protein coding genes with a 10kb window. Functional consequences for SNPs were obtained by mapping the SNPs on their chromosomal position and reference alleles to databases containing known functional annotations, including ANNOVAR, Combined Annotation Dependent Depletion (CADD), RegulomeDB (RDB), and chromatin states (only brain tissues/cell types were selected). Next eQTL mapping was performed on significant (FDR q < 0.05) SNP-gene pairs, mapping to GTEx v7 brain tissue, RNAseq data from the CommonMind Consortium and the BRAINEAC database. Chromatin interaction mapping was performed using built-in chromatin interaction data from the dorsolateral prefrontal cortex, hippocampus and neuronal progenitor cell line. We used a FDR q < 1×10^−5^ to define significant interactions, based on previous recommendations, modified to account for the differences in cell lines used here. SNPs were also checked for previously reported phenotypic associations in published GWAS listed in the NHGRI-EBI catalog.

### Gene-based and gene set analysis with MAGMA

Gene-based analysis was performed with the FUMA implementation of MAGMA. SNPs were mapped to 18,222 protein coding genes. For each gene, its association with PTSD was determined as the weighted mean χ^2^ test statistic of SNPs mapped to the gene, where LD patterns were calculated using ancestry appropriate 1KGP reference genotypes. Significance of genes was set at a Bonferroni-corrected threshold of p = 0.05/18,222 = 2.7×10^−6^.

To see if specific biological pathways were implicated in PTSD, gene-based test statistics were used to perform a competitive set-based analysis of 10,894 pre-defined curated gene sets and GO terms obtained from MsigDB using MAGMA. Significance of pathways was set at a Bonferroni-corrected threshold of p = 0.05/10,894 = 4.6×10^−6^. To test if tissue-specific gene expression was associated with PTSD, gene set-based analysis was also used with expression data from GTEx v7 RNA-seq and BrainSpan RNA-seq, where the expression of genes within specific tissues were used to define the gene properties used in the gene-set analysis model.

### Functional follow-up of the African ancestry top hit rs115539978

#### Cell Culture Experiments, RNA extraction and qPCR

Lymphoblastoid cell lines (LCLs) from the AFR superpopulation were obtained from the Coriell Institute, NJ (**Supplementary Table 6**, N = 6 lines each for the homozygous major and homozygous minor allele). Cells were cultured in RPMI 1640 medium with GlutaMAX (Thermo Scientific, 61870-036) supplemented with 15% FBS (Thermo Scientific, 26140079) and 1X Antibiotic-Antimycotic (Thermo Scientific, 15240-062) at 37C and 5% CO2 in a humidified incubator. For Dexamethasone (Dex) treatment, a final concentration of 100nM Dex (Sigma Aldrich) in 100% Ethanol was added to the medium for a total of 4 hrs. All experiments were run in duplicates.

RNA was extracted using the Quick-RNA MiniPrep Kit (Zymo, R2060) according to the instructions of the manufacturer including an additional DNase digestion. RNA concentrations were quantified via Qubit and cDNA was generated using the SuperScript IV First Strand Kit (Life Technologies, 18091200) according to the manufacturers instructions. SYBR green qPCR reactions were carried out in duplicates using POWERUP SYBR Green Master Mix (Life Technologies, A25743) and published^244^ or custom primer pairs (**Supplementary Table 7**) according to the manufacturers recommendations. Data were analyzed using the ΔΔCt method^245^ and GAPDH as reference. Between group differences were calculated using one-way ANCOVA with sex as covariate. Significance threshold was set at p = 0.05.

### Deep phenotyping exploration of the African ancestry top hit rs115539978

#### Neuroimaging

Scanning of 87 GTPC subjects took place on a 3.0 T Siemens Trio with echo-planar imaging (Siemens, Malvern, PA). High-resolution T1-weighted anatomical scans were collected using a 3D MP-RAGE sequence, with 176 contiguous 1 mm sagittal slices (TR/TE/TI = 2000/3.02/900 ms, 1 mm^3^ voxel size). T1 images were processed in Freesurfer version 5.3 (https://surfer.nmr.mgh.harvard.edu). Gray matter volume from subcortical structures was extracted through automated segmentation, and data quality checks were performed following the ENIGMA 2 protocol (http://enigma.ini.usc.edu/protocols/imaging-protocols/), a method designed to standardize quality control procedures across laboratories to facilitate replication. Briefly, segmented T1 images were visually examined for errors, and summary statistics and a summary of outliers ± 3 *SD* from the mean were generated from the segmentation of the left and right amygdala and hippocampus. Regional volumes that were visually confirmed to contain a segmentation error were discarded.

#### Startle Physiology

The physiological data of 299 GTPC subjects were acquired using Biopac MP150 for Windows (Biopac Systems, Inc., Aero Camino, CA). The acquired data were filtered, rectified, and smoothed using MindWare software (MindWare Technologies, Ltd., Gahanna, OH) and exported for statistical analyses. Startle data were collected by recording the eyeblink muscle contraction using the electromyography (EMG) module of the Biopac system. The startle response was recorded with two Ag/AgCl electrodes; one was placed on the orbicularis oculi muscle below the pupil and the other 1cm lateral to the first electrode. A common ground electrode was placed on the palm. Impedance levels were less than 6 kilo-ohms for each participant. The startle probe was a 108-dB(A)SPL, 40 ms burst of broadband noise delivered through headphones (Maico, TDH-39-P). The maximum amplitude of the eyeblink muscle contraction 20-200 ms after presentation of the startle probe was used as a measure of startle magnitude.

### Estimating PTSD heritability (*h^2^_SNP_*)

SNP-based heritability estimates (*h^2^_SNP_*) in EUA subjects were calculated using meta-analysis data including summary-level data based on LD-score regression (LDSC). Estimates are calculated for the combined PGC freeze 2 samples and separately for PGC1.5 (without UKBB), the UK biobank (including alternative subject/phenotype selections), and for men and women. Unconstrained regression intercepts were used to allow for potential inclusion of related subjects and residual population stratification, and precomputed LD scores from 1KGP EUR populations were used. For population prevalence, conservative low (10%), moderate (30%) and very high (50%) population prevalences were assumed, as reported for subjects after trauma exposure (^246^), and sample prevalence was set to the actual proportion of cases in the data. The proportion of inflation of test statistics that was due to polygenic signal rather than other causes was estimated as 1 - (LDSC intercept −1)/(mean observed chi-square −1).

To compare *h^2^_SNP_* across different ancestries, an unweighted linear mixed ^247^ was used, as implemented in the LDAK software.^248^ For each ancestry group (EUA and AFA, respectively), imputed individual-level genotype data was filtered to bi-allelic SNPs with MAF ≥ 1% in the corresponding 1KGP phase 3 superpopulation. Imputed genotype probabilities ≥0.8 were converted to best-guess genotype calls, and for each ancestry group, studies were merged and SNPs with <95% genotyping rate or MAF <10% removed. Second, to estimate relatedness between subjects, a genetic relatedness matrix (GRM) was constructed based on autosomal SNPs that were LD pruned at r^2^>0.2 over a 1MB window, and an unweighted model with α = −1, where α is the power parameter controlling the relationship between heritability and MAF. To prevent bias of *h^2^_SNP_* due to cryptic relatedness, strict relatedness filters were applied. For pairs with relatedness values > the negative of the smallest observed kinship (-0.014 for EUA and −0.045 for AFA, respectively), one subject was randomly removed. PC’s were then calculated in the remaining set of unrelated subjects. Finally, to estimate *h^2^_SNP_*, an unweighted GRM was estimated without LD-pruning, and *h^2^_SNP_* was calculated on the liability scale using REML in LDAK, including 5 PCs and dummy indicator variables for study (number of studies - 1) as covariates.

### Polygenic scoring

Polygenic risk scores (PRS) were calculated in hold out samples based on SNP effect sizes from the UKBB PTSD GWAS and publicly released major depressive disorder (MDD) GWAS (excluding 23andme; https://www.med.unc.edu/pgc/results-and-downloads). The larger female subset of the UKBB was used as a discovery set to calculate PRS in UKBB male subset. The overall UKBB was used as a discovery set to calculate PRS in PGC1.5 and PGC1.5 female subset. The overall MDD was used as a discovery set to calculate PRS in the female subset of PGC 1.5. Summary statistics were filtered to common (MAF > 5%), well imputed variants (INFO > 0.9). We also removed indels, ambiguous SNPs, and variants in the extended MHC region (chr6:25-34 Mb). LD pruning was performed using the --clump procedure in PLINK1.9, where variants were pruned if they were nearby (within 500 kb) and in LD (r^2^ > 0.3) with the leading variant (lowest p-value) in a given region. PRS were calculated in PRSice v2.1.2 using the best guess genotype data of target samples, where for each SNP the risk score was estimated as the natural log of the odds ratio multiplied by number of copies of the risk allele. PRS was estimated as the sum of risk scores over all SNPs. PRS were generated at multiple p-value thresholds (pT) (0.001, 0.01, 0.05, 0.1, 0.2, 0.3, 0.4, 0.5, 1). PRS were used to predict PTSD status under logistic regression, adjusting for 5 PCs and dummy study indicator variables, using the glm function in R 3.2.1. Plots of PRS were based on quintiles of PRS, where odds ratios were calculated in reference to the lowest quintile. For these plots, we used PRS calculated at pT=0.3 (pT=0.2 for MDD), as that was the approximate threshold that explained the most variance (most significant threshold) across analyses. The proportion of variance explained by PRS was estimated as the difference in Nagelkerke’s R^2^ between a model including PRS plus covariates and a model with only covariates. R^2^ was converted to the liability scale assuming a 30% prevalence, using the equation found in Lee et al.^249^ P-values for PRS were derived from a likelihood ratio test comparing the two models.

### Genetic correlations of PTSD with other traits and disorders

Bivariate LD score regression (LDSC) was used to calculate pairwise genetic correlation (*r_g_*) between PTSD and 235 traits with publically available GWAS summary statistics on LD Hub.^15^ Summary statistics for PTSD studies were restricted to the EUA meta-analysis, including UKBB subjects (23,212 cases, 151,447 controls) and significance was evaluated based on a conservative Bonferroni correction for 235 phenotypes (i.e. correlated traits and traits measured twice in independent studies were counted independently).

In addition, these phenotypes were compared with genetic correlations reported for PTSD and several psychiatric disorders, including 221 phenotypes and MDD^22^, 172 phenotypes and Schizophrenia (SCZ)^13^, 196 phenotypes and bipolar disorder (BPD),^23^ and 219 phenotypes and attention-deficit/hyperactivity disorder (ADHD).^24^

### Conditional analyses of PTSD top hits to test for disease specific effects

To evaluate if effects of top variants were specific to PTSD, we conditioned PTSD on MDD, and MDD plus BPD plus SCZ using the multi-trait conditional and joint analysis (mtCOJO) ^27^ feature in GCTA. MDD was selected here as the main psychiatric trait because of the high co-morbidity and genetic correlation of depressive symptoms and PTSD (*r_g_* = 0.80 for depressive symptoms and *r_g_* = 0.62 for MDD; see **Supplementary Table 15**). Publicly available summary statistics were supplied as program inputs: “Bipolar cases vs. controls”^250^ for BPD, and “MDD2 excluding 23andMe”^251^ for MDD (both from https://www.med.unc.edu/pgc/results-and-downloads); “Schizophrenia: CLOZUK+PGC2 metaanalysis”^252^ for SCZ (http://walters.psycm.cf.ac.uk/). The effect of each psychiatric disorder on PTSD was estimated using a generalized summary-data based Mendelian randomization analysis of significant LD independent psychiatric trait SNPs (r2 < 0.05, based on 1000G Phase 3 CEU samples), where the threshold for significance was set to p < 5 × 10^−7^ due to having less than the required 10 significant independent SNPs at the program default p < 5 × 10^−8^ for MDD. Estimates of heritability, genetic correlation, and sample overlap of psychiatric trait and PTSD GWAS were estimated using pre-computed LD scores based on 1000G Europeans that were supplied with LDSC (https://data.broadinstitute.org/alkesgroup/LDSCORE/eur_w_ld_chr.tar.bz2).

## Acknowledgments

This work was funded by NIMH/U.S. Army Medical Research and Materiel Command Grant R01MH106595 to CMN, IL, KJR, and KCK, and supported by 5U01MH109539. Statistical Analysis were carried out on the NL Genetic Cluster computer (URL) hosted by SURFsara. Genotyping of samples was provided in part through the Stanley Center for Psychiatric Genetics at the Broad Institute and the Cohen Veterans Bioscience. This research has been conducted using the UK biobank resource under application number 16577.

This work would have not been possible without the financial support provided by Stanley Center for Psychiatric Genetics at the Broad Institute, One Mind, and Cohen Veterans Bioscience. We are very grateful to the investigators who comprise the PGC-PTSD working group, and especially the more than 206,000 research participants worldwide who shared their life experiences and biological samples with PGC-PTSD investigators. Full acknowledgements are in the **Supplementary Note**.

## Author Contributions

### PGC-PTSD management group

M.H., K.C.K., I.L., C.M.N., A.C.P., K.J.R.

### Writing group

E.G.A., C.-Y.C., K.W.C., J.R.I.C., S.D., L.E.D., T.K., K.C.K., M.W.L., A.X.M., C.M.N., A. Ratanatharathorn, K.J.R., M.B.S., K.T.

### Study PI or co-PI

A.E.A., A.B.A., S.B. Andersen, P.A.A., S.B. Austin, E.A., D.B., D.G.B., J.C.B., L.J.B., J.I.B., A.D.B., B.B., G.B., J.R.C., M.J.D., J.D., D.L.D., K.D., A.D.-K., C.R.E., L.A.F., N.C.F., B.G., J.G., E.G., C.G., A.G.U., M.A.H., A.C.H., D.M.H., M. Jakovljevic, I.J., T.J., K.-I.K., M.L.K., R.C.K., N.A.K., K.C.K., H.R.K., W.S.K., B.R.L., I.L., M.J.L., C.M., N.G.M., M.R.M., R.E.M., K.A.M., S.A.M., D. Mehta, W.P.M., M.W.M., C.P.M., O.M., P.B.M, B.M.N., E.C.N., C.M.N., M.N., S.B.N., A.L.P., R.H.P., M.A.P., K.J.R., V.B.R., P.R.B., K.R., S.E.S., S.S., J.S.S., A.K.S., J.W.S., S.R.S., D.J.S., M.B.S., R.J.U., E.V., J.V., Z.W., T.W., D.E.W., C.W., R.Y., R.M.Y, H.Z., L.A.Z.

### Obtained funding for studies

A.B.A., P.A.A., S.B. Austin, J.C.B., L.J.B., A.D.B., B.B., G.B., J.D., C.R.E., N.C.F., J.D.F., C.E.F., E.G., C.G., M.H., R.H., M.A.H., A.C.H., D.M.H., M. Jett, E.O.J., T.J., K.-I.K., N.A.K., K.C.K., W.S.K., B.R.L., I.L., M.J.L., C.M., N.G.M., R.E.M., K.A.M., S.A.M., W.P.M., M.W.M., C.P.M., O.M., P.B.M, E.C.N., C.M.N., M.N., M.A.P., K.J.R., B.O.R., A.K.S., S.R.S., M.H.T., R.J.U., E.V., J.V., Z.W., T.W., D. E.W., R.M.Y, L.A.Z.

### Clinical

P.A.A., E.A., D.B., D.G.B., J.C.B., L.J.B., E.A.B., A.D.B., M.B., A.C.B., J.R.C., M.F.D., S.G.D., A.D.-K., C.R.E., N.C.F., J.D.F., C.E.F., S.G., E.G., A.G.U., G.G., R.H., D.M.H., M. Jakovljevic, E.O.J., A.G.J., K.-I.K., M.L.K., A.K., N.A.K., N.K., W.S.K., B.R.L., L.A.M.L., C.E.L., M.J.L., J.M.-K., D. Maurer, S.A.M., S.M., P.B.M, H.K.O., M.S.P., E.S.P., A.L.P., M.P., M.A.P., A.O.R., B.O.R., A. Rung, A.V.S., J.S.S., C.M.S., J.A.S., M.H.T., W.K.T., E.T., M.U., L.L.V.D.H., E.V., Y.W., Z.W., T.W., M.A.W., D.E.W., S.W., E.J.W., J.D.W., K.A.Y., L.A.Z.

### Contributed data

O.A.A., P.A.A., D.G.B., L.J.B., A.D.B., M.B., R.A.B., J.R.C., J.M.C.-D.-A., A.M.D., D.L.D., C.R.E., A.E., N.C.F., D.F., C.E.F., S.G., B.G., S.M.J.H., D.M.H., I.J., A.G.J., M.L.K., R.C.K., A.P.K., W.S.K., L.A.M.L., C.E.L., I.L., B.L., M.J.L., M.R.M., A.M., K.A.M., O.M., P.B.M, C.M.N., M.N., S.B.N., M.O., M.S.P., E.S.P., A.L.P., M.P., M.A.P., K.J.R., V.B.R., P.R.B., A. Rung, S.E.S., S.S., J.S.S., D. Silove, S.R.S., M.B.S., E.T., L.L.V.D.H., M.V.H., T.W., D.E.W., S.W., J.D.W., R.Y., K.A.Y., L.A.Z.

### Statistical analysis

L.M.A., A.E.A.-K., E.G.A., C.-Y.C., J.R.I.C., S.G.D., L.E.D., M.E.G., B.G., S.D.G., G.G., X-J.Q., M.W.L., A.L, A.X.M., C.M., A.R.M., S.M., D. Mehta, R.A.M., C.M.N., M.P., J.P.R., S.R., A.L.R., N.L.S., D.Schaven, C.M.S., N.S.

### Bioinformatics

L.M.A., A.E.A.-K., E.G.A., M.P.B., C.-Y.C., J.R.I.C., N.P.D, S.G.D., M.E.G., G.G., S.D.L., A.L, A.X.M., A.R.M., D. Mehta, D. Schijven, N.S., C.W.

### Genomics

M.B.-H., M.P.B., J.B.-G., K.D., M.H., S.H., M.A.H., T.K., S.D.L., J.J.L., A.C.P., K.J.R., B.P.F.R., D. Schijven, C.H.V., D.E.W.

